# Confidence judgements from fused multisensory percepts

**DOI:** 10.1101/2025.09.05.674507

**Authors:** Nicola Domenici, Pascal Mamassian

**Affiliations:** Laboratoire des systèmes perceptifs, Département d’études cognitives, École normale supérieure, PSL University, CNRS, 75005 Paris, France; Applied Cognitive Psychology, Ulm University, Germany

**Keywords:** Metacognition, Metaperception, Multisensory Integration, Temporal Bisection, Modelling

## Abstract

Distinguishing between reliable and unreliable internal sensory representations is crucial for a successful interaction with the environment. Precise estimates of the uncertainty of sensory representations are critical to optimally integrate multiple sensory modalities and produce a coherent interpretation of the world. However, once a multisensory percept is produced, it is not yet known whether humans still have access to the uncertainty of each sensory modality. Here, we asked human participants to perform a series of temporal bisection tasks, in either unimodal visual or auditory conditions, or in bimodal audiovisual conditions where vision and audition could be congruent or not. The validity of their temporal bisection was assessed by asking participants to choose which of two consecutive decisions they felt more confident of being correct, focussing only on the visual modality. We found that once multisensory information is integrated, participants could no longer access the unisensory information to evaluate the validity of their decisions. Comparing three generative models of confidence, we show that confidence judgments are fooled by the fused bimodal percept. These results highlight that some critical information is lost between perceptual and metaperceptual stages of processing in the human brain.

## Introduction

Trusting our senses is a crucial step to safely navigating any environment: perceiving up to the most subtle event is not as useful if one cannot consistently evaluate whether it is safe to plan, adapt, and act based on that very information. Creating accurate and precise internal representations of the environment is thus necessary, but not sufficient, to achieve optimal interactions with the external world. To this end, humans also need to assess the reliability of these perceptual decisions. Overall, the ability to self-monitor and self-control one own’s perception is defined as metaperception^1^, and is part of the broader concept of metacognition^2^ which includes general knowledge about any given mental state^3^.

To date, metaperception has been investigated mostly considering one sensory modality at a time^1,4–11^. When more than one sensory modality has been tackled, it was mostly to compare confidence sensitivities and biases across modalities^12–14^. More recently, confidence judgments were measured for a couple of multisensory illusions, such as the sound-flash illusion and the McGurk effect, where the same percept can be obtained from different combinations of sensory information. Results were somewhat inconsistent in that confidence was either larger when the multisensory cues were congruent than incongruent^15^, or similar in these two conditions^16^. These studies indicate that participants are sometimes able to detect when sensory cues are incongruent, possibly because the multisensory percepts are not perfect metamers and retain some unisensory properties.

Faced with multiple sources of information that can be slightly incongruent, the brain continuously integrates and segregates information from different senses, often merging two (or more) cues into coherent, unitary percepts^17,18^. Through such a process, known as multisensory integration, perceptual cues are optimally combined according to their reliability to minimize the variance of the final estimate^17^. Thus, a noisy signal will show reduced influence in determining the final, multimodal percept, as it will be weighted less than a more reliable cue^19^. But are sensory cues necessarily lost following integration? In an elegant study, Hillis and colleagues^20^ addressed this question in two conditions, when the two cues belong to a single sensory modality (here, stereopsis and texture for the visual perception of 3D slant), and when the two cues belong to different senses (here, vision and touch). When combining two cues within a single sensory modality, participants could no longer access individual cues to discriminate stimuli differing across one single dimension, indicating that the two sensory cues were forcefully fused. However, when the cues came from different senses, participants were able to use a combination of mandatory fusion and Single-cue estimation, suggesting that unisensory information was still accessible even though that cue was integrated with another sensory modality.

Considering the results of Hillis and colleagues, here we asked whether once integrated into a fused object, unisensory cues can still be accessed to inform subsequent metaperceptual judgments. To address this question, we developed an ensemble of temporal bisection tasks using different combinations of visual, auditory, and audiovisual stimulations. Within such a design, participants were asked to report whether the second stimulus of a sequence of three was closer in time to the first or the third. Metaperceptual performance was then evaluated by using a Confidence Forced-Choice task^21,22^, in which participants had to choose between two temporal bisection decisions the one they believed was more likely to be correct. While confidence ratings are often involved in similar studies (being intuitive and relatively faster), referring to a confidence forced-choice overcomes the subjective usage of rating scales which can fluctuate within and between participants^21^. Furthermore, confidence ratings can sometimes be misinterpreted to indicate a more refined sensory decision when simultaneously providing perceptual and confidence responses^21,23^. In order to avoid confusions for the participants and erroneous interpretations of the results, we always asked participants to base their perceptual decisions and confidence judgments on the visual cue alone (when available). By doing so, we were able to evaluate whether the final confidence judgment originated from that visual information alone, or was tied to the integrated bimodal percept.

To evaluate metaperceptual performance, we adapted an established confidence generative framework^22^ and contrasted three different models. The Single-cue model assumed that metaperceptual evidence was only computed from the unisensory information that was relevant, in our task the visual one, irrespective of multisensory integration. The comparative model assumed that metaperceptual evidence was computed independently from both visual and auditory modalities, and then merged at a subsequent step. Lastly, the integrative model assumed that metaperceptual evidence was computed directly from the optimally integrated audiovisual percept, thus being inextricable from the multisensory integration processes.

## Results

We collected data from 15 participants in a series of temporal bisection tasks^24^, in which triads of visual and/or auditory stimuli (S1, S2, and S3) were sequentially presented (Figure 1A). Via manipulation of the delay between S1 and S2, temporal performance was evaluated by asking participants to indicate whether S2 was closer in time to S1 or S3. Participants performed five different temporal bisection conditions: visual-only (V), auditory-only (A), audiovisual synchronous (AVs, in which auditory and visual stimuli were presented at the same time), and two audiovisual asynchronous (AV+ and AV-, in which the auditory component of S2 was shifted compared to the visual one by +100 and -100ms, respectively). In the first phase of the experiment (perceptual phase), we measured perceptual performance in all five bisection conditions to evaluate its suitability in eliciting optimal audiovisual integration. A sample psychometric function for a given participant (S01) is shown in Figure 1B, together with the four stimulus difficulty levels (delays between S1 and S2) that were then chosen for the rest of the study. These difficulty levels were selected individually for each participant to lead to probabilities of 0.65 or 0.85 to report that S2 was closer to S1 or S3. In the second phase of the experiment (metaperceptual phase), we measured metaperceptual abilities using a Confidence Forced-Choice paradigm (Figure 1C). In this paradigm, for each trial, participants had to make a perceptual decision on a first triad of stimuli to be bisected, followed by another bisection decision shortly afterward on a second triad. Then, once both decisions were collected, participants were asked to choose which decision they felt more likely to be correct. Similarly to the perceptual phase, participants performed five different conditions (V, A, AVs, AV+, and AV-). For the three bimodal conditions, the first bisection decision was performed on a visual sequence of stimuli, while the second one on an audiovisual sequence.

**Figure 1.**
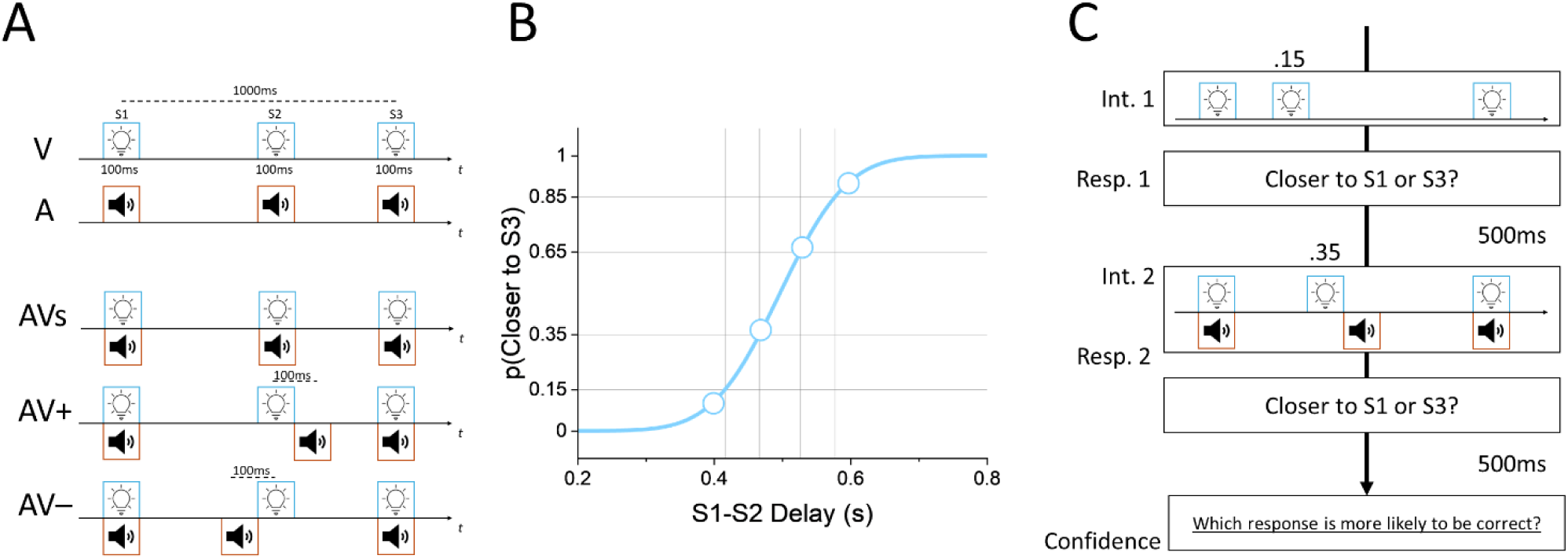
A) Temporal bisection for the perceptual task, grouped per condition. B) Psychometric curve obtained from participant S01 in the visual perceptual task. Data collected from four interleaved staircases are grouped in four equally sized bins (blue circles, each one made up of 30 trials) for visualization. From the psychometric curve fitted to the raw data (solid blue curve), we sampled four points (S1-S2 delays corresponding to 0.15, 0.35, 0.65, or 0.85 probabilities to respond that S2 was closer to S3) that were subsequently used to generate confidence pairs for the metacognitive task. C) Sample confidence trial for the metacognitive task (using the AV+ condition for illustration purposes; note how the first decision was made on a visual sequence of stimuli). Participants performed a first temporal judgment (Interval 1), followed by another temporal bisection (Interval 2), before reporting which decision (Response 1 or Response 2) they felt more likely to be correct (Confidence).

### Perceptual Phase

To gauge perceptual biases and sensitivities, we fitted a psychometric function to each participant’s bisection decisions. From the psychometric function, we estimated the bisection point (BP; i.e. the S1-S2 delay for which S2 was equally likely to be judged closer to S1 or S3) and just noticeable difference (JND; i.e. the increase in S1-S2 delay to reach 75% correct) values for each participant in each experimental condition. For this analysis, no participants were excluded.

From the literature on multisensory integration, we expect a reduction of the JND in all the audio-visual conditions, and a shift of the BP in the audio-visual conflict conditions^17^. To test the significance of the shift of the BP in the AV+ and AV– conditions, we ran a one-way repeated measure ANOVA with Condition (V/A/AVs/AV+/AV–/AV+_pred_/AV–_pred_) as the only factor, including in the analysis the shift for both AV+ and AV-conditions as predicted by multisensory optimal integration. As expected from multisensory fusion, we found that BP significantly changed with condition (F(6,84)=43.57, p-value<0.001, η^2^=0.76, BF_10_ = 9.17·10^21^). In particular, we found that shifts in BP aligned with the predictions of optimal integration, significantly decreasing in the AV+ condition and increasing in the AV– one (Figure 2; Supplementary Information: Statistical Analyses). To test the significance of the reduction of the JND in the audio-visual conditions, we ran a one-way repeated measure ANOVA with Condition (V/A/AVs/AV+/AV–/optimal) as the only factor. Our results highlighted the significance of the Condition factor (after applying the Greenhouse-Geisser correction due to the sphericity assumption being violated: F(5,70) = 11.979, p<0.001, η^2^ = 0.461, BF_10_ = 5.52 * 10^5^). Post-hoc analyses confirmed that the JNDs for the AV conditions were significantly lower than both unisensory JNDs, and did not differ from the values predicted by optimal multisensory integration, (Figure 2; Supplementary Information: Statistical Analyses).

Taken together, these results confirmed that participants optimally integrated multisensory information in all three bimodal conditions^17^. Assessing consistent multisensory integration in our temporal bisection task was a key prerequisite in developing a proper design to investigate multisensory confidence (something we did not satisfactorily achieve in previous attempts to address this issue^25^). Having certified this step, we proceeded with the same perceptual task to test how confidence evidence was accumulated following multisensory integration.

**Figure 2.**
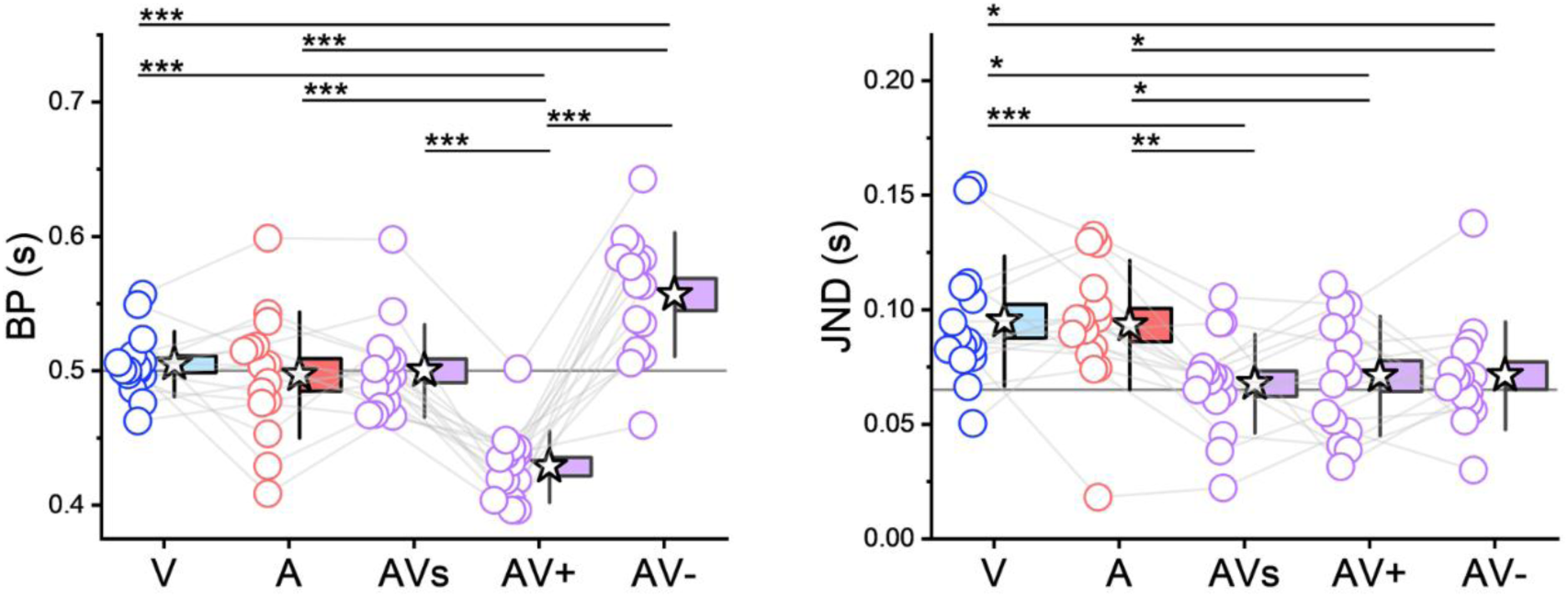
Comparisons across conditions for both the Bisection Point (BP, left panel) and Just Noticeable Difference (JND, right panel). For the BP, the horizontal line indicates the physical bisection point, while for the JND it indicates the optimal bimodal sensory noise as predicted by the Maximum-Likelihood Estimate. Circles represent individual participants, while stars denote group means. Box-plots extend up to ±SEM, and error bars indicate ±SD (* = p-value<0.05, ** = p-value<0.01; *** = p-value<0.001).

### Metaperceptual Phase

#### Unisensory Metacognitive Performance

Before testing our models on all bimodal conditions, we first need to ascertain that participants were able to re-evaluate their perceptual performance when only one sensory modality was present. This was crucial, since all parameters extracted from the unisensory conditions were then subsequently used as fixed parameters in the Single-cue, Comparative, and Integrative models. To increase the efficiency of parameters estimation, for any given condition we performed the analysis on the aggregated data (so that for each condition we recover parameters from 2,880 confidence trials). Pooling data from all participants was a sensible strategy, since confidence pairs were created using stimuli with comparable difficulty between participants, thus controlling for inter-individual variability (see Figure 1B and Supplementary Information: Experimental Procedure). For analytical purposes, S1-S2 delay values for the different expected performance were extracted from the aggregated psychometric functions obtained in the corresponding sensory task. We extracted values from the unisensory Visual and the Auditory conditions, and generated predictions for the bimodal conditions starting with the Visual stimulus and adding an auditory cue to S2 that was either synchronous (AVs), followed (AV+) or preceded (AV–) the visual cue by 100ms (see both Methods and Supplementary Information: Experimental Procedure, Figure S1).

All parameters were estimated considering a previously developed generative confidence model^22^. This metacognitive framework emphasizes that confidence can be impaired by some confidence noise, but can also benefit from some confidence boost, that is information available at the confidence inference stage that was not available for perceptual decision. Following this framework, we extracted four parameters for each sensory modality: the sensory criterion θ (how biased was the perceptual decision), the sensory noise σ (how sensitive was the perceptual decision), the confidence boost α (what was the fraction of new evidence that was used for confidence that was not available for the perceptual decision), and the confidence noise 𝜀 (how uncertain was the confidence judgment). These parameters were estimated by considering all the different confidence comparisons (Figure 3). We divide these conditions in four groups of choice probabilities between the two intervals, where each interval can take any of the four difficulty levels. Each group corresponds to a particular pair of perceptual decisions in the two intervals (e.g. responding that S2 was closer to S1 in the first interval, and closer to S3 in the second interval; see top-left panels in Figure 3A and 3B). For each group, the probability of choosing the first decision as the most confident one is plotted as a function of the S1-S2 delay for the first (x-axis) and second temporal decisions (colored lines). Assuming that all perceptual decisions and confidence pairs are mutually independent, best model parameters were then obtained by maximizing the summed log-likelihood of each confidence pair. Best-fitting values are reported in Table T1, while continuous colored lines indicate model’s predictions for the corresponding stimuli combination. Dot sizes are proportional to the number of trials for that particular combination of stimulus difficulties and perceptual responses.

**Figure 3.**
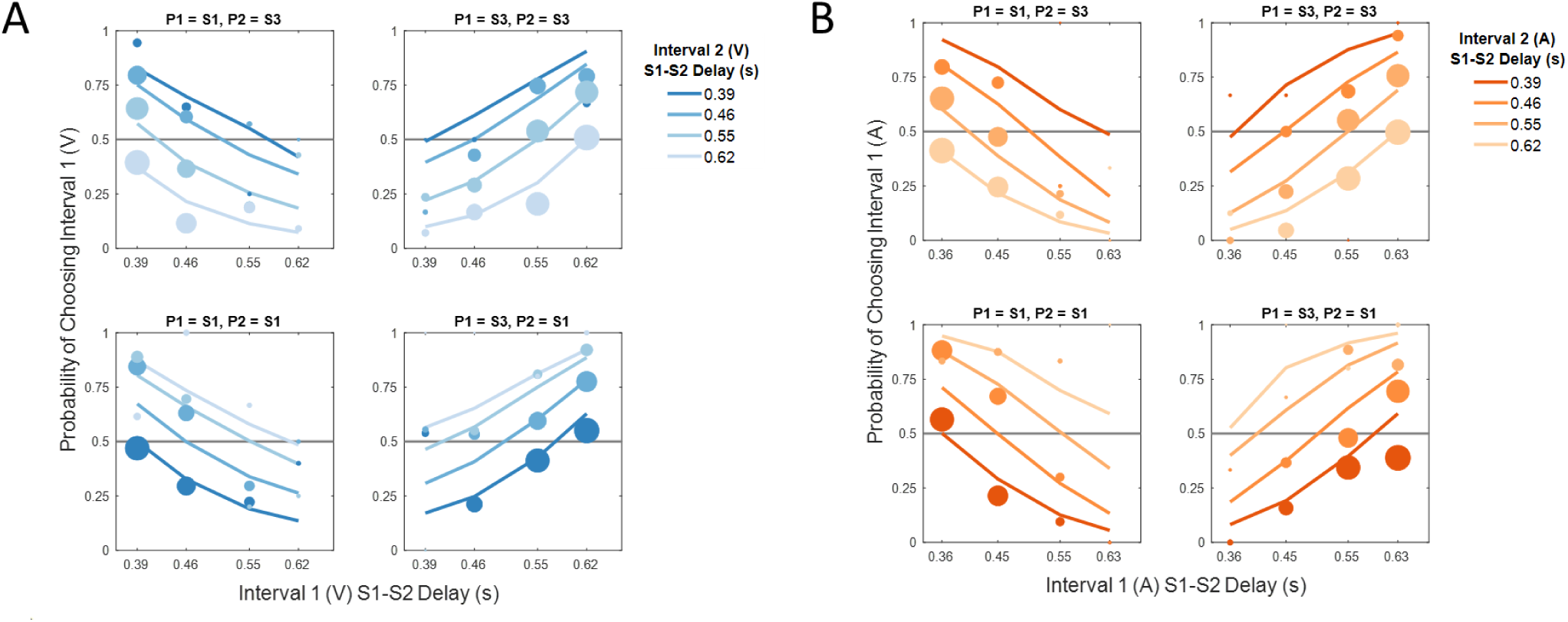
Unisensory metacognitive performance and model fits. A) Groups of confidence probabilities to judge that the perceptual decision in interval 1 was more likely to be correct than that in interval 2 for the purely visual stimuli. Each panel shows one particular group of perceptual decisions for each interval (P1, P2, as reported above each scatter plot). Stimulus difficulties for the interval 1 varies along the x-axis, and stimulus difficulties for the interval 2 are shown by different blue color saturations. Size of the dots is proportional to the number of trials for each combination of stimulus difficulties and perceptual decision. Colored lines are predictions of the best fitted model. B) Same as A) for the auditory confidence judgments.

We found clear signs of metaperceptual capabilities when considering how well participants were able to distinguish between correct and incorrect decisions (Visual confidence efficiency: 0.959; Auditory confidence efficiency: 0.618; see ^22^ for the definition of confidence efficiency). To appreciate how good the confidence judgments are, we can take for example the group of visual decisions where S2 was reported closer to S3 in both intervals (top-right panel of Figure 3A). When stimuli in both intervals had an S1-S2 delay of 0.62 s, both perceptual decisions were mostly correct, and because the stimulus difficulty was matched across the two intervals, participants were equally confident to choose the first or second interval. As the S1-S2 delay for the first interval was shortened (from 0.62 to 0.39 s, from right to left on the x-axis), if participants kept reporting that S2 was closer to S3, they made more and more mistakes. Critically, this increase of mistakes in the first interval is associated with a decrease of confidence choices in favor of that interval. Overall, confidence judgments were closely monitoring perceptual performance, a bit better for vision than for audition in our temporal bisection task.

**Table T1.** Best fitting parameters for both unisensory metacognitive tasks.

| | $\theta$ | $\sigma$ | $\alpha$ | $\varepsilon$ |
| --- | --- | --- | --- | --- |
| V | 0.483 | 0.079 | 0 | 0.183 |
| A | 0.476 | 0.069 | 0.959 | 1.654 |

#### Learning effects and adjustments of perceptual parameters

The bimodal conditions were run in blocks to avoid some plausible instability of the weights assigned to each modality that would occur from the random jumps across audiovisual conditions. Because of this choice of blocking conditions, that was following the same order for all participants, we need to consider some possible learning effect that could affect our results. We did find some small shifts in the perceptual biases and in the bimodal sensitivities. Because it is critical for the confidence analysis that we know precisely how difficult a perceptual decision was to the participants, we have adjusted the perceptual bias and sensitivity before fitting any model. Specifically, we accounted for the constant shifting between visual and audiovisual decisions by introducing a switch cost on the JND for the audiovisual estimates. In addition, we allowed for a possible recalibration due to audiovisual temporal asynchronies by introducing a small updating of the unisensory BPs (See Supplementary Information: Adjustments of Perceptual Biases and Sensitivities).

#### Multimodal Models of Metaperception

In order to reveal the source of the metacognitive evidence that was used in the audiovisual confidence judgments, we designed and tested three different models (Figure 4): the Single-cue, the Comparative, and the Integrative model (for a more detailed explanation of the models’ theoretical implications, see Supplementary Information: Metacognitive Models).

The Single-cue model assumes that, regardless of audiovisual integration, all information involved in metacognitive computations comes solely from the visual stimulus. This model can be seen as the ideal one, since participants were explicitly asked to evaluate their visual performance. According to this model, internal evidence used to draw confidence judgments can be enhanced by information drawn from the original visual stimulus (in the form of visual confidence boost), and can be corrupted by confidence noise that is visual in nature. Overall, the Single-cue model depicts a strategy where participants used the visual modality as the computing focal point to generate their confidence decision, disregarding additional information from the auditory modality.

**Figure 4.**
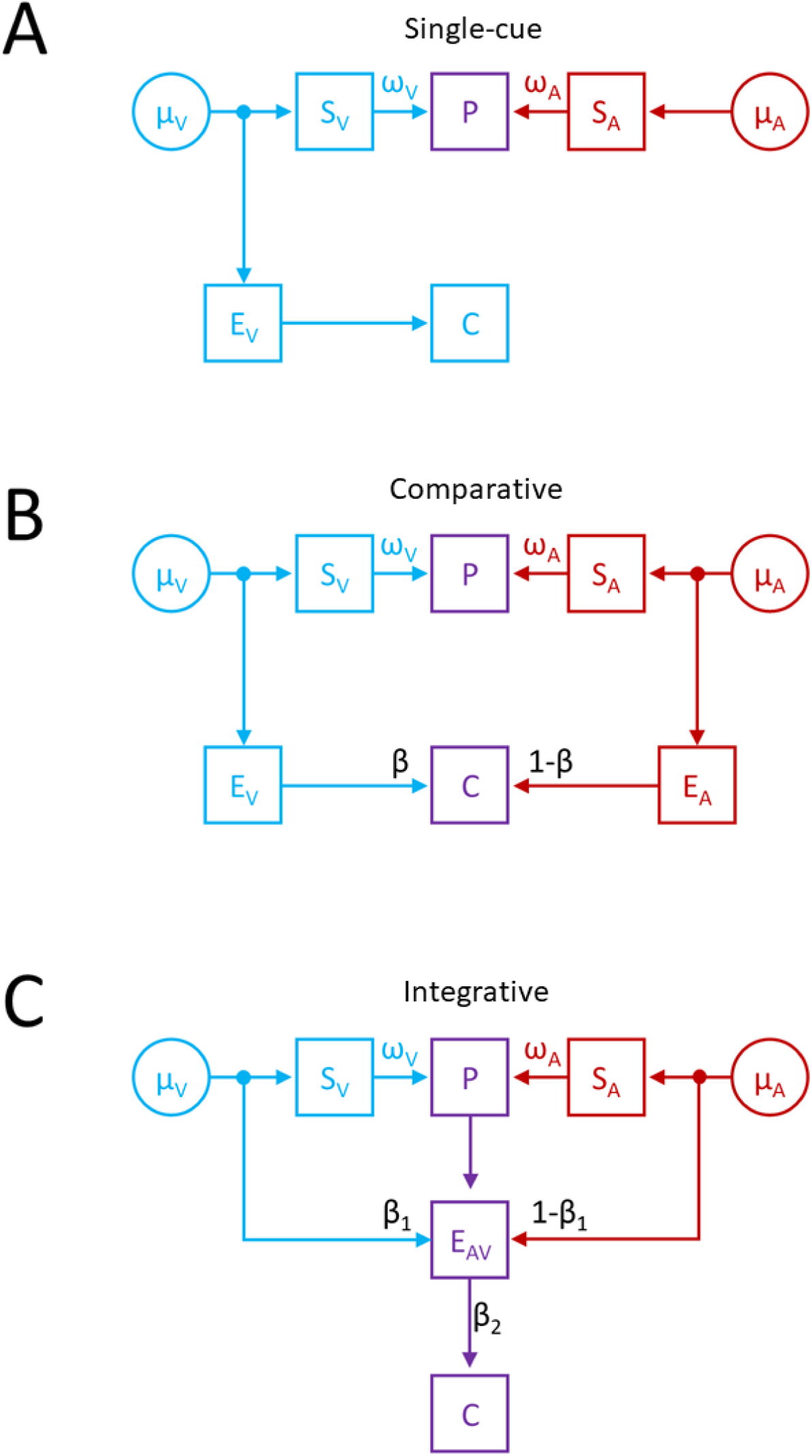
Visualization of the Single-cue (A), Comparative (B), and Integrative (C) models. At the sensory stage, all models implied optimal multisensory integration, suggesting sensory estimates were combined into audiovisual percepts (P) according to the corresponding weighting factors 𝜔_*V*_ and 𝜔_*A*_. At the metaperceptual stage, however, the three models depicted significantly different behaviors. The Single-cue model assumed that the confidence evidence C was generated from the visual modality alone. The Comparative model assumed that confidence evidence was generated independently from each individual sensory cue, and then combined at a later stage according to a factor *β*. Lastly, the Integrative model assumed that all confidence computations originated from the optimally integrated bimodal percept. Consequently, C was enhanced by a combination of unisensory confidence boosts (weighted according to *β*_1_) and corrupted by audiovisual confidence noise, modelled as a rescaled sum of unisensory confidence noises (via the rescaling factor *β*_2_).

The Comparative model, on the other hand, describes a situation where auditory and visual modalities independently contribute to the generation of metaperceptual judgments. Specifically, the model assumes that confidence evidence is individually generated from each sensory stimulation, and only subsequently is merged at a later processing stage. According to the Comparative model, visual and auditory information is first enhanced and corrupted separately by visual and auditory confidence boost and noise. Then, the resulting visual and auditory confidence information is combined by a weighting factor *β* (which was left free to vary), to generate the audiovisual confidence evidence.

Lastly, the Integrative model assumes that all confidence computations originate from the audiovisual, optimally integrated percept. Therefore, any additional information used to refine the confidence evidence, identified by the confidence boost, is forcefully derived from the bimodal estimate. To represent this interaction, we combined and rescaled unisensory confidence boosts by a weighting factor *β*_1_. Following similar logic, the internal confidence evidence is corrupted by some confidence noise which, in this model, is a combination of unisensory confidence noises. We thus considered the bimodal confidence noise to be equal to the sum of the visual and auditory confidence noise, rescaled by a factor *β*_2_. In this model, both *β*_1_ and *β*_2_ were left free to vary.

#### Multisensory Metacognitive Performance

For all bimodal conditions, parameter estimation was conducted similarly to what was already done for each unisensory metacognitive condition. We emphasize that all values for the models’ fixed parameters were extracted from both the unisensory V and A metacognitive conditions (see Table T1). To compare our three models, we pooled empirical data in sets of confidence choices groups. Then, for each condition and each model, we simulated 1,000 participants, subsequently collapsing them into one dataset to obtain one probability per confidence choice represented in the scatter plots. Assuming independence across perceptual decisions for all confidence pairs, and independence across sensory modalities from each other, we then obtained sets of best model parameters by summing the log-likelihood of each pair, comparing participants metaperceptual performance with the performance simulated from each model. To account for the different number of degrees of freedom between models, we resorted to the Aikake Information Criterion (AIC), which is based on the summed log-likelihood with a penalty factor for increased degrees of freedom. Results of the comparison between models are shown in Figure 5A. Overall, the model that best captured participants’ performance was the Integrative one, while the Comparative followed second by a significant margin.

Best-fitting values for the free parameters are worth mentioning. For the Comparative model, we found that best-fitting *β*=0, while for the Integrative models we obtain as best *β*_1_=0.53 and *β*_2_=0.59, respectively. The final value of *β* for the Comparative model is intriguing because, being zero, it suggests that the confidence evidence for the bimodal stimuli comes solely from the auditory modality, even though the task of the participants was to report their confidence about their visual decision. This remark further reduces the plausibility that the Comparative model is the most appropriate. The interpretation of the parameters of the Integrative model is more revealing. As a reminder, in the Integrative model, *β*_1_ represents the normalization factor for the confidence boost, and *β*_2_ the coefficient scaling the sum of confidence noises. Intuitively, we initially thought that the normalization factor *β*_1_ defining multimodal confidence boost would have followed both sensory weights (𝜔_*V*_ and 𝜔_*A*_) in our sample. Indeed, if audiovisual estimates are the results of optimal integration, and each individual cue influences the final percept according to their corresponding 𝜔, we hypothesized that confidence boost rescaling should undergo similar dynamics and maintain congruent ratios. In line with this prediction, *β*_1_ was relatively close to the 𝜔_*V*_ observed in the perceptual task (which was 0.43, as can be verified from values reported in Table T1 and equation S6 in the Supplementary Information: Metacognitive Models section) and, consequently, (1 − *β*_1_) was relatively close to 𝜔_*A*_ (especially considering how sensory fits were modulated to account for the switches between visual and audiovisual decisions; See Supplementary Information: Adjustments of Perceptual Biases and Sensitivities). Regarding *β*_2_, while its value is lower than 1, thereby suggesting that bimodal confidence noise stems from the potential combination of unisensory confidence noises, we need to remain cautious. Since confidence noises were very different across modalities, we cannot rule out that overall bimodal confidence noise was mostly determined by the worse metacognitive modality. Further studies will be helpful to shed light on the nature of this interaction.

**Figure 5.**
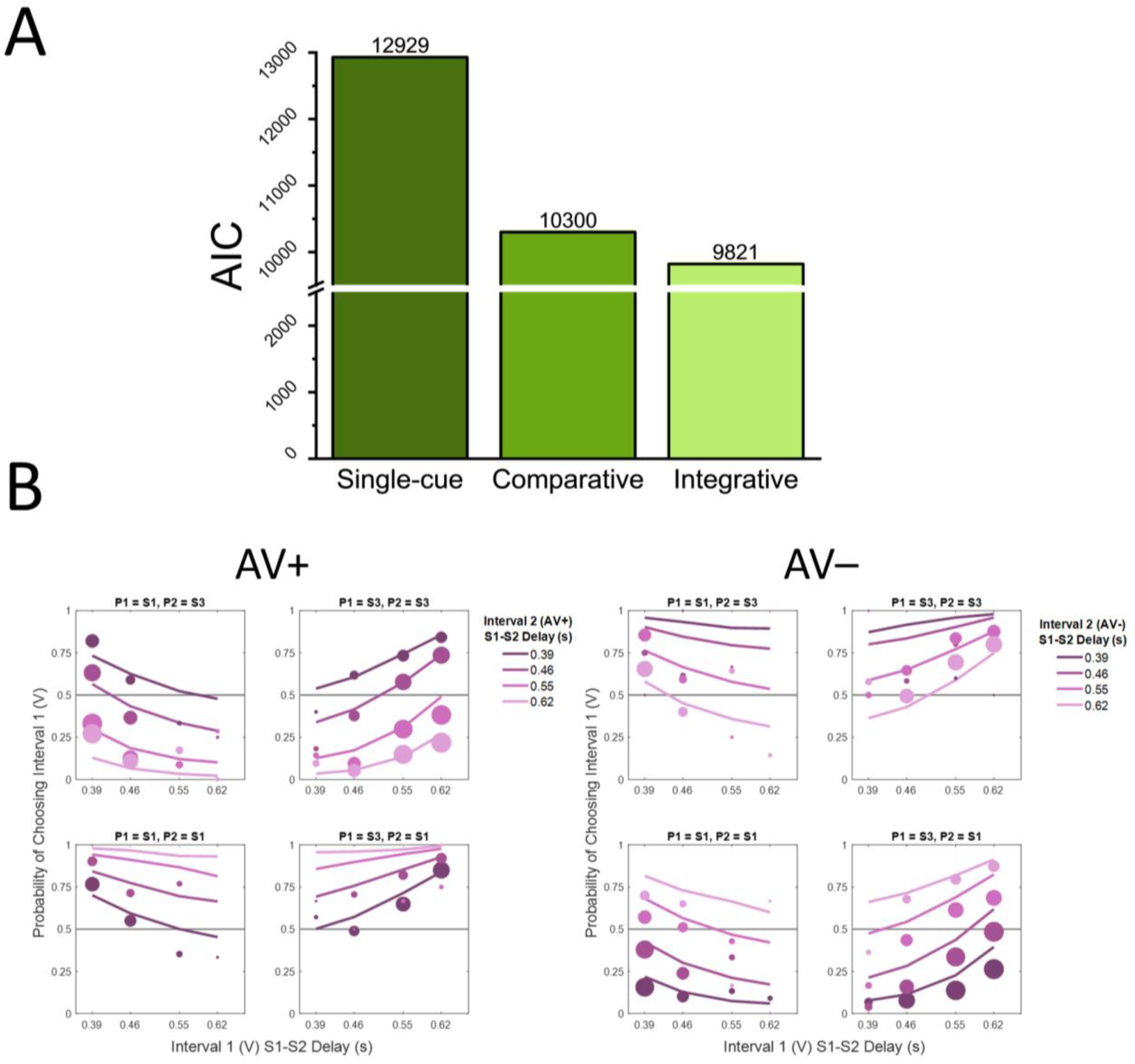
Model comparison for the audiovisual metaperception, and best fits of two audiovisual conditions. A) Comparison between the three models using the AIC estimator (smaller is better). The Integrative model was a better predictor of multisensory confidence judgments than the other two models. B) Metaperceptual performance for the two audiovisual cue conflict conditions (the third audiovisual condition is shown in Supplementary Information: Figure S5). The left panels show the group of confidence probabilities for the AV+ condition, and the right panels for the AV– condition. Scatter plots follow the same formats as those used in Figure 3. Note that in these conditions, the first interval always contained a vision only stimulus (x-axis) and the second interval an audiovisual stimulus (different color saturations). Continuous lines correspond to the performance of the Integrative, best-fitting model.

To further expand on the Integrative models’ behavior, we plotted all model confidence choices on top of humans’ probabilities for both AV+ and AV– conditions (Figure 5B, colored lines). Due to their relevance for our study, here we focused on the two audiovisual conditions that presented participants with conflicting sensory information (similar results for the AVs condition are reported in the Supplementary Information, section: Goodness-of-fit for the Confidence Models). Overall, the Integrative model adequately replicated human’s confidence choices across different configurations of stimulus strengths, audiovisual sensory conflicts, and perceptual decisions, suggesting that not only it was the best one at replicating human behavior, but also that its performance did not significantly deviate from human data. Additional comparisons between models and empirical data can be found in the Supplementary Information, section: Goodness-of-fit for the Confidence Models.

## Discussion

In the current study, we investigated how humans build the confidence they have about their correctness in a visual task, in a context where an auditory cue was optimally integrated with the visual stimulus to bias the final percept and make it more reliable. Using a series of temporal bisection tasks, we modelled confidence using a previously developed generative framework, specifically testing three different models. The Single-cue model assumed that humans can selectively access unisensory cues to generate confidence evidence, regardless of the sensory integration occurring at the perceptual level. Alternatively, the Comparative model assumed that confidence is independently computed for both sensory estimates, and then combined in a subsequent step to produce a multimodal confidence judgment. Finally, the Integrative model assumed that confidence evidence is forcefully generated from the integrated percept, plausibly occurring at a later processing stage.

The confidence judgments in our experimental conditions were best accounted for by the Integrative model, suggesting that multimodal confidence evidence stems from the bimodal percept and cannot be backtracked to individual cues. Intriguingly, Hillis and colleagues^20^ argued that unisensory signals can still support additional perceptual comparisons, even after being integrated with cues from other modalities. This crucial finding indicates that perceptual estimates are not lost following multisensory integration and thus appears in contradiction with our results.

It may be argued that the difference between the study of Hillis et al.^20^ and ours is the use of different sensory modalities and perceptual tasks, but we doubt that this is the case for two reasons. First, multisensory integration in humans have been shown to follow similar mathematical rules regardless of the senses and domains in play^26^. Second, metaperceptual abilities appear very similar across sensory modalities^12–14^ and tasks^27^. In sum, we have no reason to believe that the differences in sensory modalities and perceptual tasks are responsible for the apparent discrepancy between the study of Hillis and colleagues and ours.

The apparent contradiction between the original study^20^ and ours may stem from a conflation of two different levels of processing. These two levels have been discussed since the early days of metacognition^28^ and correspond here to perceptual and metaperceptual processes. There is evidence that these two levels have different corresponding neural bases. At the perceptual level, neural correlates of multisensory integration have been found in primary sensory cortices by resorting to a range of techniques, such as blood-oxygenation level-dependent (BOLD) signal amplitudes analysis^29^, event-related potentials (ERPs) comparison^30^, and magneto-encephalography (MEG) source reconstruction^31^. These findings highlight the role of early sensory cortices in multimodal processing, which are actually multisensory in function^32,33^, and suggest that the integration of multimodal signals is localized at early information processing stages. At the metaperceptual level, neural correlates of metacognitive computations are likely to involve prefrontal cortical regions, and thus later processing stages. For instance, using functional magnetic resonance imaging (fMRI), Fleming and colleagues found evidence of the involvement of the right rostrolateral prefrontal cortex when providing confidence ratings during a perceptual categorization task^34^. Other studies highlighted the role of additional prefrontal cortical regions in modulating metacognition, such as the right inferior frontal gyrus^35,36^, the anterior cingulate cortex^37,38^, and the orbitofrontal cortex^39^.

In summary, we argue that the two levels of processing for perception and metacognition, occurring in distinct brain regions, is the source of the (apparent) discrepancy between the results of Hillis and colleagues and ours. When performing multimodal sensory comparisons, unisensory cues are still accessible because they are encoded in analogous cortical regions. However, once a commitment to a perceptual decision is made and metacognitive evidence is engaged, the sensory information at the origin of the confidence judgment is the optimally integrated percept. From a phylogenetic standpoint, there might have been evolutionary pressure to rely on a combination of single cue estimators and fusion to generate perceptual decisions, since sensory cues might or might not come from the same source, even when sharing similar spatial coordinates^16,20^. Conversely, computing confidence judgments from evidence other than the one related to the final perceptual decision is most often disadvantageous: once the brain commits to one decision, the subsequent logical step is to evaluate the reliability of that decision before exploring possible alternatives. One exception might be error monitoring, where the metacognitive process of detecting an error in our perceptual decision would probably benefit from an access to the unisensory cues to determine, for instance, whether one cue was very discrepant from the others. Future work should look at the benefits and costs of separating perceptual and metaperceptual processing stages at the neural level. But such a separation would explain the supramodal nature of metaperception^40^ that we discussed above, where humans can easily make comparative metacognitive judgments not only within^27^, but also between sensory modalities^12–14,41^. This view is consistent with the proposal that metacognition is domain-general in that it is not tied to individual sensory modalities, nor to specific features within a modality^42^.

On a final note, it is worth discussing the peculiarities of the Integrative model and the resulting values of the best-fitting parameters. Other than representing the best model to account for our data, it also showed interesting benefits coming from its implementation. First, we found that the bimodal confidence boost follows a similar weighted combination across senses as the one at the multisensory fusion stage. This result further supports the validity of the Integrative model, suggesting that audiovisual information used both for the perceptual decisions and metacognitive judgments are strongly related. This is also intuitively unsurprising, considering that the confidence boost determines the amount of information used for the metaperceptual judgment that was not used for the perceptual decision: if one sensory modality drives the final percept more than the other, it should also similarly affect multimodal metaperception. Second, our analysis suggested that multimodal confidence noise might just be the sum of the unisensory confidence noise. However, given the unisensory confidence noise values reported in Table T1, caution should be taken as auditory confidence noise was significantly higher than the visual one. Future studies should attempt to find conditions where confidence noises for auditory and visual judgments are similar in order to test the optimality of the metacognitive integration.

To conclude, we showed that multimodal confidence judgments are strongly tied to the integrated multisensory percept and cannot be inferred solely from individual sensory cues, even when participants are explicitly asked to directly focus on a single cue. Following previous neural studies focusing on either multisensory integration or metacognition, our results provide behavioral evidence that metaperceptual computations take place at a later processing stage than multisensory integration. In addition, once metaperceptual processing starts, individual sensory cues are no longer accessible.

## Materials and Methods

### Participants

A total of 15 participants took part in the study (mean age: 27.13 ± 5.23y, 3 Males, 12 Females) after being recruited via the French RISC database (http://www.risc.cnrs.fr). All participants had normal or corrected-to-normal vision and normal hearing. Data collection was performed at the École Normale Supérieure, in Paris, in a dark, small room suited to control sound reverberation. Testing procedures were performed in accordance with the Declaration of Helsinki and approved by the local ethics committee (Comité d’Ethique de la Recherche de l’Université Paris Cité), and all participants provided signed informed consent before taking part in the experiment. Participants were compensated for their participation (10 euros/hour, for a total of 30 euros).

### Exclusion Criteria

To test whether psychometric curves for the perceptual task were reliable, we evaluated for each nonlinear fit the corresponding R^2^. Since all nonlinear fits were statistically significant, and all R^2^>0.8 (Mean: 0.93, SD: 0.04; Range: [0.8 0.99], we concluded that no psychometric function failed to capture participant’s performance in the sensory task.

At the participant’s level, the only exclusion criterion we applied involved the JND. Within the current study, S2 was presented in a temporal window between 0.2 and 0.8 seconds (from S1 and considering the overall S1-S3 duration of 1 second). Given the physical bisection point at 0.5 s, any participant with a JND larger than 0.3 second (i.e. the distance between 0.5 and either 0.2 or 0.8) would have been excluded due to the impossibility of correctly evaluating temporal precision. No participants showed this behavior, so we included all of them in the analysis.

### Apparatus and Stimuli

All experiment scripts were generated using Matlab 2020b and the Psychtoolbox routine (v. 3.0.1). Visual stimuli were displayed on a CRT monitor (Vision Mater Pro 454) with a resolution of 800 x 600 pixels and a refresh rate of 100 Hz, with participants placing their head on a chin rest at a viewing distance of 57 cm. Auditory stimuli were delivered using a soundbar (Bose Solo SoundBar Series 2, black color) plugged via an aux cable to an external USB audio card (Audioengine D3 DAC Audio USB).

Visual stimuli were white disks with a diameter of 3° of visual angle, centered 5° below a central fixation point and displayed on a gray background for 100ms. No temporal smoothing was applied to the boundaries of the visual stimulation. Auditory stimuli were 110ms bursts of pink noise characterized by a 10ms cosine ramp at both the beginning and end (thus considered both the onset and offset of the auditory stimuli at the midpoint of the cosine ramp). The vertical distance between the visual stimulus and the center of the speaker was 15° of visual angle. Stimuli’s effective durations and synchronization were monitored by measuring them via Tektronix TBS series oscilloscope before data collection.

### Experimental Procedure

In all conditions, three stimuli (S1, S2, and S3) were delivered in succession, interleaved with variable delays, and the participants’ goal was to indicate whether S2 was closer in time to S1 or S3. The total duration of the stimulus to be bisected (S1-S3) was always fixed at 1000 ms, regardless of the sensory modalities involved. In all audiovisual bisection tasks, participants were always asked to respond considering only the visual cue.

### Perceptual Task

The whole study was divided into two separate phases: in the first phase, sensory performance for all possible temporal bisection tasks was evaluated. For each task, participants completed 120 trials in which the delay between S1 and S2 was defined on a trial-by-trial basis by a staircase adaptive procedure to accelerate stochastic approximation^43^. Four staircases were simultaneously run (30 trials per staircase), with two of them converging at 25% probability to indicate that S2 was closer in time to S3, and the other two converging at 75% probability. The order of presentation of all conditions was identical for all participants, so that the visual task was the first one to be completed, followed by the auditory one, the audiovisual synchronous, and the two audiovisual asynchronous performed last (with the AV+ task performed before the AV–).

### Metaperceptual Task

The second phase of the experiment was focused on evaluating metaperceptual performance, using a Confidence Forced-Choice design^22^. In such a task (Figure 1C), observers perform a first perceptual decision (in our study, a first temporal bisection), followed by a comparable, second one. Once the two responses are independently provided, participants must report which decision they believe is more likely to be correct (hence, the forced choice).

### Fitting Procedures

For the perceptual task, we evaluated performance by fitting psychometric functions into each individual participant’s data (Figure 1B). For all conditions, the resulting curve represents the proportion of responses ‘closer to S3’ expressed as a function of the delay between S1 and S2. Data were fitted using a cumulative Gaussian, and the best fit approximating empirical data was obtained by minimizing the negative log-likelihood. As a result, each fitting procedure allowed the definition of two crucial parameters: the bisection point (BP), which is the delay between S1 and S2 being perceived as equal to the delay between S2 and S3, and the Just Noticeable Difference (JND), which represents the increase in S1-S2 delay needed so that participants can consistently distinguish whether S2 is closer to S1 or S3 75% of the time. While the BP indicates overall temporal bias–thus, the closer to the physical bisection the more accurate the performance–the JND is a parameter used to represent temporal sensitivity. As a general rule, the lower the JND the better the performance.

### Statistical analyses

To compare sensory performance between conditions, we use both inferential and Bayesian parametrical analyses in the form of repeated-measure ANOVAs. For frequentist analyses, statistical significance was discussed assuming an alpha level of 0.05, and post hoc comparisons were performed adjusting for multiple hypotheses testing using Bonferroni correction. For the Bayesian analyses, BF_10_ for main effects were reported as a comparison to the null model. Posterior odds for post hoc comparisons were corrected for multiple testing by fixing to 0.5 the prior probability that the null hypothesis holds across all comparisons^44^. For all analyses, outliers were defined considering values that fell outside the ±3 SD range.

### Models and Parameters Estimation

To understand how metaperceptual evidence was accumulated from the audiovisual percept, we referred to an already existing confidence generative framework^22^, aiming at capturing participants’ behavior in all bimodal conditions: AVs, AV+, and AV–. The three models presented in the current study were developed by adapting the original conceptualization. Key parameters of the original generative model of interest to the current work are explained below.

The confidence noise (𝜀) identifies the reduction in sensitivity to the confidence evidence (similarly to the sensory noise, which reduces sensitivity to the sensory evidence); it is assumed to be normally distributed and independent from the sensory noise. Confidence boost (α), on the other hand, includes some information involved in the confidence judgment that was not considered for the sensory decision. For instance, good decisions are often performed quickly, so if participants were able to evaluate their own reaction times, this information could be used to improve the confidence judgment. Both confidence noise and boost can also be used to estimate the confidence efficiency, which is computed as the inverse of the equivalent confidence noise variance (assuming α = 1). Confidence efficiency ranges from 0 (no metacognitive efficiency) to infinite (no confidence noise) and, albeit providing similar results compared to other efficiency measurements, should not be confused with the classical meta-d’/d’ approach^45^ as the latter is obtained by a ratio of standard deviations. For an exhaustive explanation of the generative model and its parameters, we suggest to the potentially interested readers to refer to the original description of the model^22^.

All parameters were extracted after pooling all participants’ data together. By ensuring that different S1-S2 delays across participants corresponded to identical levels of performance, we managed to control for inter-individual sensory differences (thus allowing us to safely merge all trials together). For instance, while the point closest to the left tail of the psychometric (as shown in Figure 1B) corresponded to different S1-S2 delays between observers, depending on the shape of the individual psychometric, it always coincided with a probability of reporting that S2 was closer to S3 15% of the time. Similarly, all models were generated at the pooled level and compared against the aggregated, corresponding datasets.

For all metacognitive analyses, S1-S2 delay values were extracted from the psychometric function fitted into aggregated data from the corresponding sensory task (akin to what already done at the individual level, as depicted in Figure 1B).

### Models Comparison

In order to compare the three models, we had to consider the different degrees of freedom characterizing each one. Notably, the Single-cue model had no free parameter (so, no minimization routine was required), the Comparative model had one (*β*), and the Integrative model two (*β*_1_ and *β*_2_). To account for such difference, we converted the summed log-likelihoods into corresponding AICs before testing whether one of them was best at describing participants’ behavior, according to the canonical formula

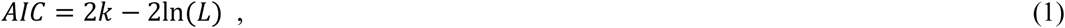

where *k* is the number of degrees of freedom, and 2ln(*L*) is the summed log-likelihood.

## Acknowledgments

This work was supported in part by the French National Research Agency (ANR) grant “VICONTE” (ANR-18-CE28-0015), Labex (ANR-10-LABX-0087 IEC), and “FrontCog” (ANR-17-EURE-0017).

## Data Availability Statement

Both raw data, source data, and MATLAB codes for the modelling part can be freely accessed at the following link: https://zenodo.org/records/15791663.

