## Supplementary Information for "Confidence judgements from fused multisensory percepts"

#### Experimental Procedure

The total duration of the interval to be bisected (S1-S3) was always fixed at 1000 ms, regardless of the sensory modalities involved. The AVs, AV+, and AV– conditions were generated by presenting an auditory stimulus either in synchrony with the visual stimulus, or 100ms before or after the visual stimulus (Figure S1).

In each Confidence Forced-choice task, all stimuli to be delivered were defined at the individual participant's level using the perceptual results from the previous phase. Specifically, for each participant, we fitted a psychometric function onto their individual data (see Main text, Fitting Procedures), and from the resulting curve, we sampled four different points (Main text, Figure 1B). These four points corresponded to the interval between S1 and S2 leading to 15, 35, 65, and 85% probability of indicating that the second stimulus was closer to S3. Notably, these four intervals were matched for difficulty at the between-subject level, denoting both easier (15% and 85%) and harder (35% and 65%) bisections so that, for instance, the shortest interval of the four always led to 15% probability of reporting that S2 was closer to S3 regardless of the participant. Intervals were also matched at the within-subject level, and balanced in difficulty depending on each S1-S2 interval's distance from the sensory criterion. Thus, the two easier bisections were equally difficult when compared to each other, just like the two harder ones.

After sampling the four intervals for each participant, to create confidence trials we paired them through all possible combinations (16), that were repeated 12 times in random order. In total, each metaperceptual condition featured 192 confidence trials. The conditions' order of presentation mirrored the one developed for the first sensory phase (V/A/AVs/AV+/AV-) and was fixed across participants to minimize, and potentially model, inter-individual switch costs.

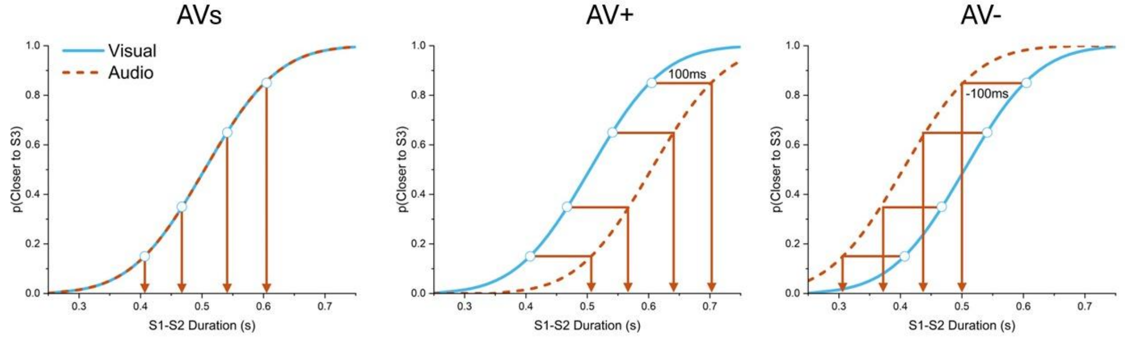


**Figure S1.** Generation routine for the bimodal temporal bisection stimuli in the metacognitive task. In all three bimodal conditions, the first temporal judgment was always performed on a visual sequence, while the second temporal bisection was always bimodal. To create bimodal intervals, we sampled four intervals from the visual psychometric obtained in the V perceptual task (blue continuous line) and then added an auditory cue to S2 depending on the condition (orange arrows). In the AVs condition (left panel), auditory cues were presented at the same time as visual ones; in the AV+ condition (middle panel), auditory cues were delivered 100ms after the visual ones; in the AV- condition (right panel), auditory cues preceded the visual ones by 100ms.

In both V and A metaperceptual conditions, confidence comparisons were performed using the four intervals extracted from the V or A perceptual condition, respectively. Specifically, confidence judgments were carried out by asking participants to compare their responses on two visual (V) or two auditory (A) bisection trials. In the three bimodal conditions, participants were instead asked to compare their response in a first, purely visual bisection task with the response given in a second, audiovisual bisection task. For both visual and audiovisual temporal bisections, all four different intervals were extracted from the psychometrics obtained in the V perceptual condition, with the only difference that in the second bisection task an auditory stimulus was paired with all visual stimuli (either synchronous, preceding, or following the second visual stimulus by 100 ms, akin to the first phase; one sample confidence trial is depicted in the main text, Figure 1C).

Each metaperceptual condition was divided into six blocks of 32 confidence pair trials, at the end of which participants were encouraged to take a small break. Overall, the procedure took three hours to complete and was divided into three sessions of one hour each. A detailed description of the order of conditions across sessions can be found in Table S1.

| Session | Task | Condition | Number of trials |
|---|---|---|---|
| Session 1 | Perceptual | V, A, AVs, AV+, AV- | 120 x 5 |
| | Metaperceptual | V | 192(confidence pairs) |
| Session 2 | Metaperceptual | A, AVs | 192 x 2 |
| Session 3 | Metaperceptual | AV+, AV- | 192 x 2 |

**Table S1.** Task and condition order across the three experimental sessions.

#### Statistical Analyses of the Perceptual Phase

We performed post-hoc analysis to detail the effect of the bisection point (BP) in the perceptual phase. After Bonferroni correction for multiple comparisons (adjusting the p-value for comparing a family of 21), we confirmed that the BP in the AV+ condition was significantly lower than the BP in the V ($t_{(14)} = 7.5$, p-value < 0.001, 95% CI [-0.108 -0.044], posterior odds: $1.93 \cdot 10^5$), A ($t_{(14)} = 6.2$, p-value < 0.001, 95% CI [-0.1 -0.036], posterior odds: 63), AVs ($t_{(14)} = 7.02$, p-value<0.001, 95% CI [-0.103 -0.039], posterior odds: $5.55 \cdot 10^3$), and AV- (t(14)=-12.6, p-value < 0.001, 95% CI [-0.16 -0.096], posterior odds: $2.66 \cdot 10^4$) ones. Conversely, the BP in the AV- condition was significantly higher than the BP in the V ($t_{(14)} = -5.1$, p-value < 0.001, 95% CI [0.02 0.084], posterior odds: 13.2), A ($t_{(14)} = -5.89$, p-value < 0.001, 95% CI [0.028 0.092], posterior odds: 460), and AVs ($t_{(14)} = -5.58$, p-value < 0.001, 95% CI [0.025 0.089], posterior odds: 89.45) ones. Notably, the BP in both the AV+ ($t_{(14)} = -2.01$, p-value = 0.996, 95% CI [-0.052 0.011], posterior odds: 0.788) and AV- ($t_{(14)} = 0.572$, p-value = 1, 95% CI [-0.026 0.038], posterior odds: 0.062) conditions did not differ from its prediction. No other comparison was statistically significant (all p-values = 1; all posterior odds < 0.074).

For the analysis of the just noticeable difference (JND), we performed post-hoc analyses (adjusting p-values for comparing a family of 15) to detail the effects across conditions. While both visual and auditory JNDs differed from the optimal fused prediction (V vs pred: $t_{(14)} = 5.6$, p-value < 0.001, posterior odds: 698.18; A vs pred: $t_{(14)} = 5.291$, p-value < 0.001, posterior odds: 6297.63), all bimodal JNDs did not (AVs vs pred: $t_{(14)} = 0.656$, p-value = 1, posterior odds: 0.126; AV+ vs pred: $t_{(14)} = 1.308$, p-value = 1, posterior odds: 0.158; AV– vs pred: $t_{(14)} = 1.091$, p-value = 1, posterior odds: 0.21). JNDs in all audiovisual conditions were significantly lower than the ones in the visual (V vs AVs: $t_{(14)} = 4.944$, p-value < 0.001, 95% CI [0.011 0.044], posterior odds: 19.266; V vs AV+: $t_{(14)} = 4.347$, p-value < 0.001, 95% CI [0.007 0.041], posterior odds: 12.789; V vs AV-: $t_{(14)} = 4.318$, p-value <0.001, 95% CI [0.007 0.044], posterior odds: 6.635) and auditory ones (A vs AVs: $t_{(14)} = 4.635$, p-value < 0.001, 95% CI [0.009 0.042], posterior odds: 122.86; A vs AV+: $t_{(14)} = 4.039$, p-value = 0.002, 95% CI [0.006 0.039], posterior odds: 3.482; A vs AV-: $t_{(14)} = 4.009$, p-value = 0.002, 95% CI [0.005 0.039], posterior odds: 7.196). All other comparisons between conditions were not statistically significant (all p-values = 1; all posterior odds < 0.07).

We report predicted BP and JND values against the empirical parameters in Figure S2.

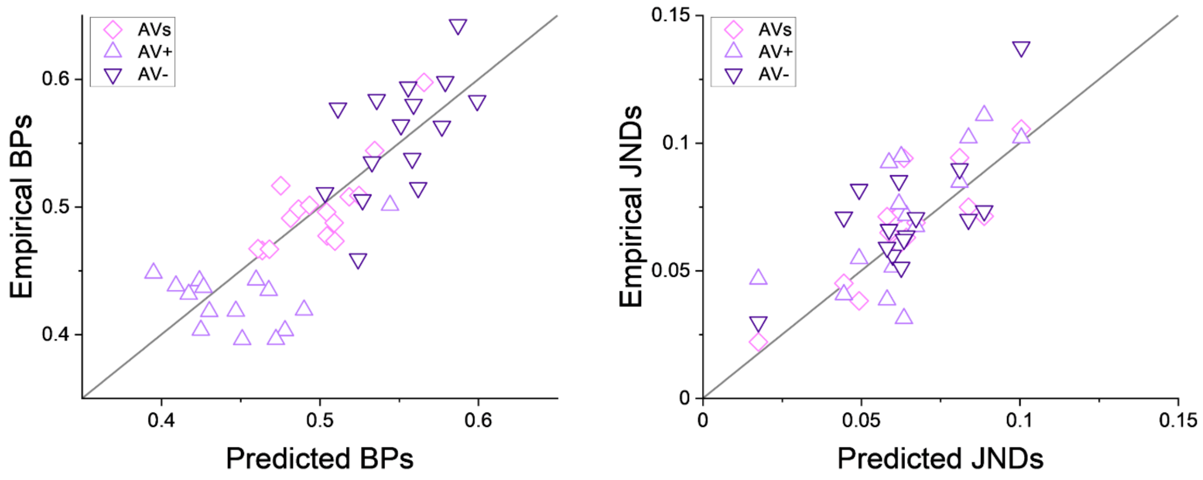


**Figure S2.** Empirical BPs (left panel) and JNDs (right panel) plotted against the predicted values from the optimal integration model.

#### Adjustments of Perceptual Biases and Sensitivities

Due to the conditions' fixed order of presentation in the metacognitive experiment, before inferring the confidence parameters for the bimodal conditions, we checked whether there was any learning effect that could compromise our results. To this end, we compared JNDs across all three bimodal conditions, independently considering perceptual decisions for visual and bimodal temporal bisections (Figure S3A). We thus performed one-way repeated measures Bayesian ANOVAs, with order of presentation as a factor and the JND as the dependent variable. Overall, for the visual bisection tasks, we found that our data were more likely under the null hypothesis rather than the alternative one including the order of conditions ($BF_{10} = 0.26$). For changes in precision considering bimodal bisections, the resulting BFs were less conclusive ($BF_{10} = 1.2$). Despite this, it is unlikely that learning effects occurred only for bimodal temporal judgments, while no change was observed for the visual ones. Thus, we excluded the possibility that the order of conditions affected temporal precisions across the experiment, and ruled out the potential influence of learning effects.

Surprisingly, when pooling all participants together, the aggregated psychometric function significantly deviated from the optimal prediction in the AV- condition (for both the visual and audiovisual temporal bisection judgement, Figure S3B; as a reminder, sensory predictions were computed simulating 1000 datasets assuming optimal integration and using parameters value extracted from the unisensory metacognitive tasks). Before interpreting this deviation, we aimed at modeling the difference between empirical and predicted psychometrics for all temporal bisections in the AVs, AV+, and AV- condition (thus, including also the performance for the first visual bisection task). Increasing the fidelity of our models at the sensory level was crucial to properly test our metacognitive hypotheses, since the latter were based on a temporally fused audio-visual event.

In order to better align the perceptual performance with the empirical one, we modelled non-specific effects affecting sensory performances (i.e., unrelated to multisensory integration and metacognition) that potentially influenced the temporal bisection task. Most notably, this procedure further helped modulate any inherent influence of the confidence-forced choice design, which required participants to constantly switch from visual to audiovisual stimuli. Thus, this modelling stage operated both at the BP and sensitivity levels, allowing us to update the unisensory bisection points (from the calibration/unisensory stages) and to introduce a sensitivity penalty (in the form of a switch cost) to the perceptual precision for the second (i.e., audiovisual) judgment.

We thus reconsidered the unisensory BPs as follows

$$\mu_V = \mu_V^{calib} + \delta_V \quad \text{(S1)}$$

$$\mu_A = \mu_A^{calib} + \delta_A \quad \text{(S2)}$$

where $\mu_V^{calib}$ and $\mu_A^{calib}$ are the visual and auditory BPs (as predicted by the unisensory conditions), respectively, and $\delta_V$ and $\delta_A$ are the corresponding empirical criterion shifts. Similarly, at the sensitivity level, we applied a switch cost to the audiovisual (but not to the visual) percept so that

$$\sigma_V = \sigma_V^{calib} \times \vartheta \quad \text{(S3)}$$

$$\sigma_A = \sigma_A^{calib} \times \vartheta \quad \text{(S4)}$$

where $\sigma_V^{calib}$ and $\sigma_A^{calib}$ are the visual and auditory JNDs (as predicted by the unisensory conditions), respectively, and $\vartheta$ is the switch cost induced by the constant shift between visual and audiovisual stimuli.

To avoid overfitting, we simultaneously perform the fitting procedure to all three bimodal conditions. We then calculated the best-fitting parameters maximizing the log-likelihood between the empirical and predicted data, obtaining the following values: $\delta_V = -0.016$, $\delta_A = 0.029$, and $\vartheta = 1.39$. In other words, relative to our expectation from the Perceptual Phase, the bisection points were shifted by less than 30 milliseconds and the just noticeable difference was impaired by about 40% in the Metaperceptual Phase. Visual comparisons between the empirical and predicted fits are reported in Figure S3B. Following our modeling of the deviation of the observed empirical psychometric functions, the biggest shifts were observed for the AV- condition, intriguingly occurring for both the visual and the bimodal temporal bisection tasks.

Although we do not have an unequivocal explanation for this, the best-fitting values obtained from the fitting procedure accounting for non-specific factors suggests that some sort of multisensory temporal recalibration processes occurred after the exposure to the previous, opposite audio-visual temporal asynchrony (the AV+ condition). Indeed, instances of audio-visual temporal recalibration have been observed after exposure to a temporal conflict, so that the perception of a subsequent asynchrony is shifted towards the exposure lag[1]. Interestingly, temporal recalibration also occurs in a stronger fashion when the first stimulus is presented through the visual modality[2], suggesting that the previous exposure to the AV+ condition might have prompted this mechanism in our study. Therefore, its presence and absence might be suitable to explain the increased noise in the bimodal bisection task. It is also noteworthy to highlight that multisensory recalibration and integration are likely two distinct processes, as integration is reliability-based[3] and recalibration is most likely based on a fixed-ratio adaptation[4]. Thus, the presence of the latter does not necessarily exclude the former.

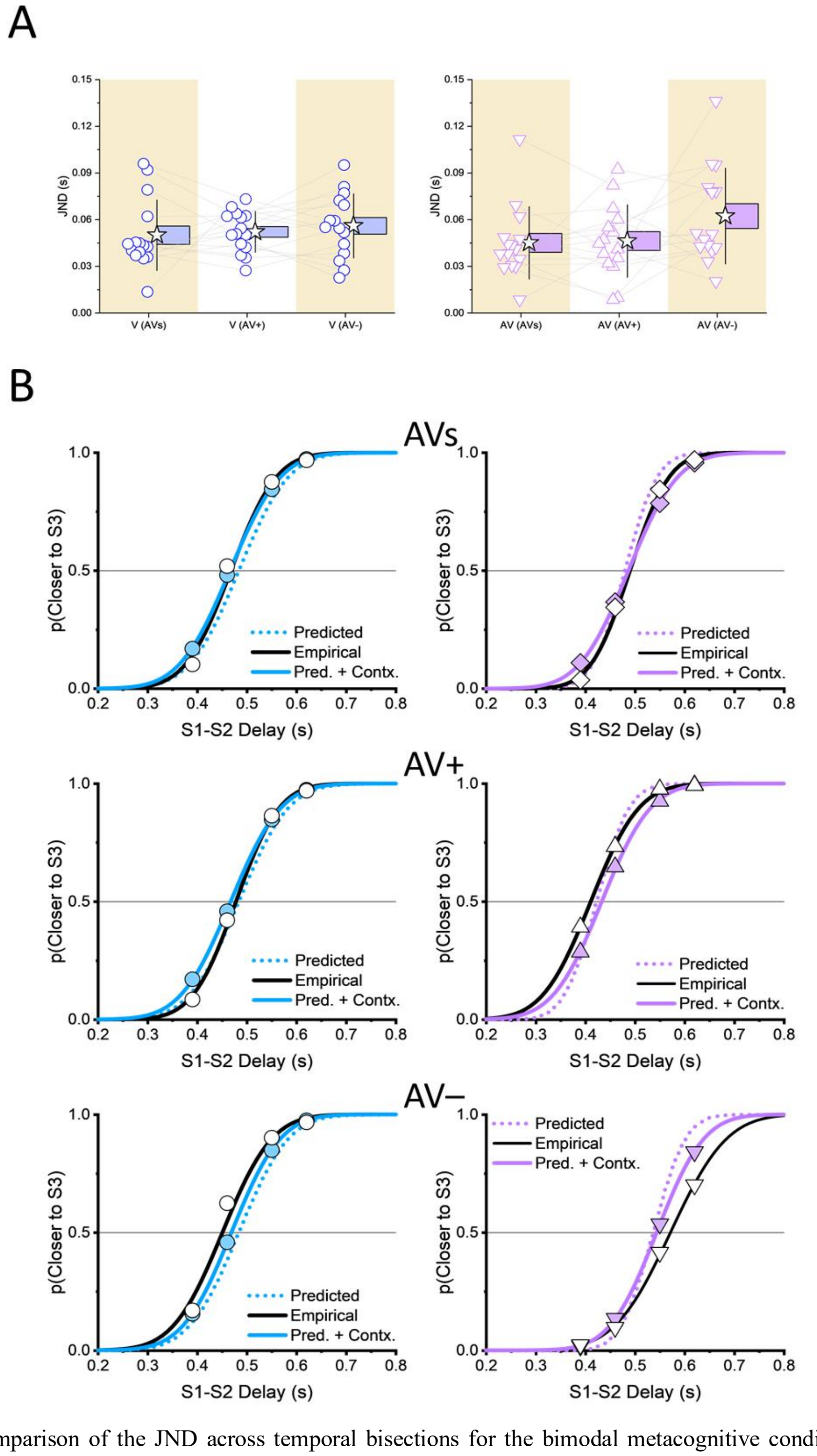


**Figure S3.** A) Comparison of the JND across temporal bisections for the bimodal metacognitive conditions. In our data, no significant change of JND was observed across the three conditions (AVs, AV+, and AV–), thus allowing us to exclude the presence of learning across conditions. B) Psychometric functions adjustments. Continuous black lines indicate empirical performance at the group level. Dotted colored lines show the predicted performance assuming optimal multisensory integration. Continuous colored lines show the adjusted psychometric functions after applying BP recalibration and JND switch cost.

Another possibility is that the previous exposure to the audiovisual asynchrony in the AV+ bisections triggered cumulative recalibration mechanism occurring during the AV- conditions. We thus checked whether the BPs changed during the session by fitting psychometric functions to individual data, dividing the performance into three equally sized blocks. We then scanned for changes due to block presentation via two-way repeated-measure Bayesian ANOVA, including as factors the Modality (V vs. A) and Block (1 vs. 2 vs. 3). Our results highlighted that the best model at approximating the data was the one including only the Modality ($BF_{10}$ = 21.49), while we also found significant evidence in favor of the null hypothesis when considering the Block factor (even when interacting with the Modality factor; all $BF_{10} < 0.15$).

Overall, we argue that the shallower psychometric curve in the AV- condition is a by-product of recalibration processes (as attested by the different signs of $\mu_V$ and $\mu_A$ in equation S1 and S2, respectively), as well as the cost, in terms of sensory precision, of constantly switching between different sensory modalities. Moreover, further artificial deflation of the slope might be influenced by how the pooled psychometric is missing its anchor to the upper bound, which was inevitable considering how S1-S2 intervals were sampled from the perceptual phase.

#### Pooled analysis and absence of idiosyncratic metacognitive strategies

While pooling participants together was possible due to how we normalized the interindividual difficulty for the task, we also had to check whether any given participant exhibited metacognitive performances that significantly differed from the rest of the group. To do so, we then resampled our dataset via Jackknife and, for each iteration, we computed the Kullback-Leibler divergence as follows

$$D_{KL} = \sum P(x) \cdot \log\left(\frac{P(x)}{Q(x)}\right) \tag{S5}$$

where $P(x)$ represents the participant probability choices, and $Q(x)$ model probability choices. Via Jackknife resampling, we obtained a distribution of 15 $D_{KL}$ values, each one obtained by removing a given participant from the entire sample. We then checked whether any specific value significantly deviated from the others by simply scanning for outliers.

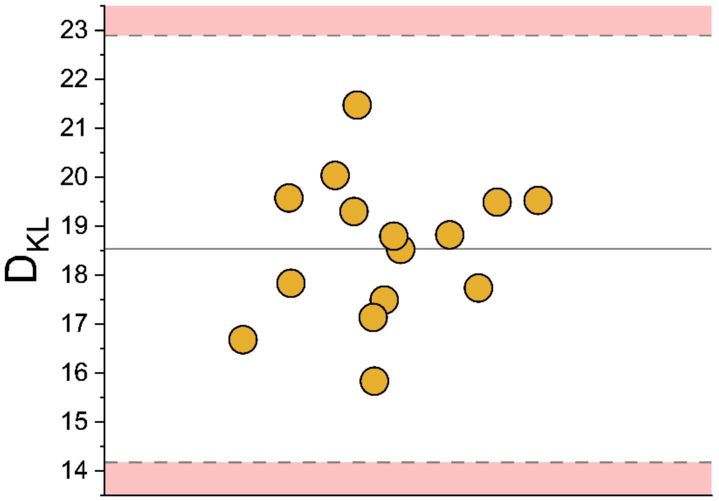


**Figure S4.** $D_{KL}$ distribution obtained via resampling, removing one participant at each iteration. To test for the absence of idiosyncratic metacognitive strategies in our sample, we resampled our dataset using the jackknife method, removing one participant at a time for every iteration. We thus checked whether among this distribution of $D_{KL}$some of the values could be categorized as outliers by falling outside the ±3 SD range. As no value fell outside such range, we concluded that participants adopted similar metacognitive strategies, which further support our pooling procedure. The continuous, grey line indicates the mean of the distribution, while the two dashed lines delimit the ± 3 SD range.

In the end, no $D_{KL}$ fell outside the ±3 SD range from the mean (Figure S4), indicating that no participant employed metacognitive strategies that differed significantly from those of the others. For reference, the mean of the $D_{KL}$ distribution was 18.54, the SD was 1.45, and the range [15.82, 21.47].

#### Goodness-of-fit for the Confidence Models

In the main text, we showed that the Integrative model best represented the human confidence probabilities compared to the Comparative and Single Cue models. Nonetheless, the procedure used (i.e., AIC) only allowed us to compare models against each other at the surface level. Here, we tested whether the Integrative model reliably approximates human confidence choices.

First, in Figure S5 we reported the Integrative model performance for the AVs condition, which was not introduced in the main text due to its reduced impact on the interpretation of the results (as no audiovisual incongruency was presented).

To test the goodness-of-fit of all three models, we fitted weighted linear regressions between the human and model confidence probability choices (see Figure S5 for the AVs condition and Figure 5B in the main text for the AV+ and AV- conditions). Weights for each point in these regressions were defined by the number of trials performed by human participants for that given combination of stimulus strengths and perceptual responses. We then computed the Root Mean Squared Error (RMSE) separately for all conditions and all models and reported their values in Table S2. As already

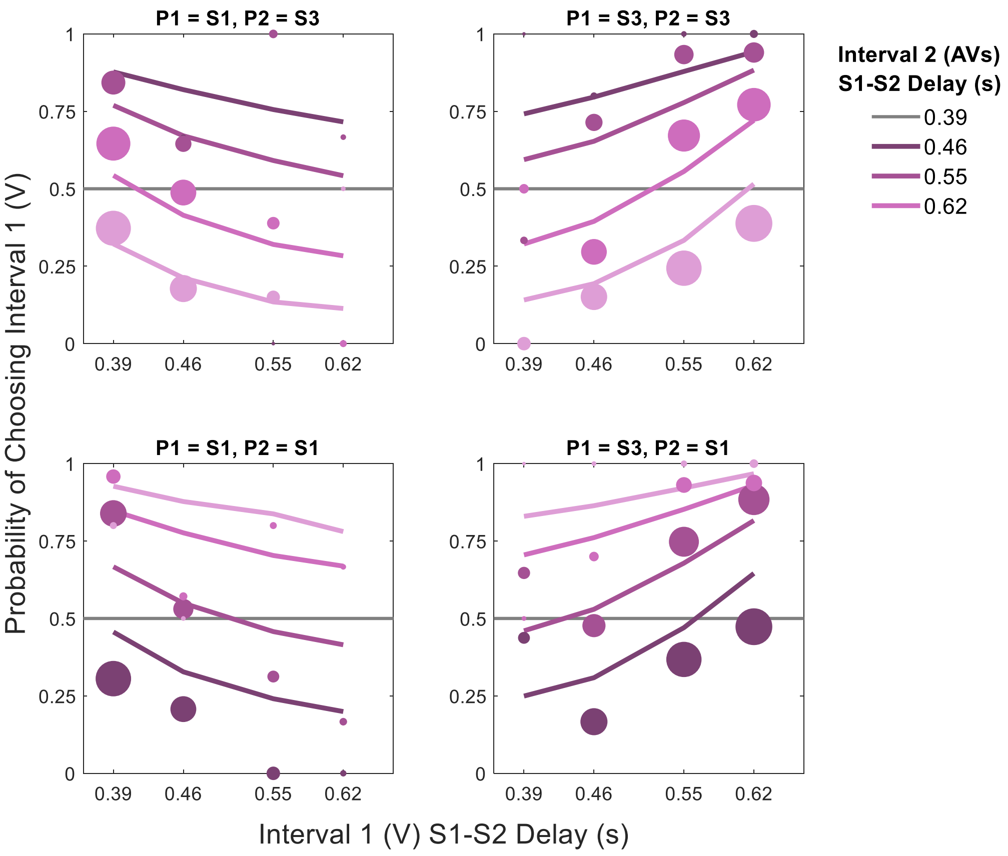


**Figure S5.** Groups of confidence probabilities for the AVs condition. Similarly to Figure 5B in the main text, scatter points report the probability of choosing the first sensory decision as the most confident one, with this probability being expressed as a function of both perceptual decisions and S2-S3 interval duration for the second bisection (indicated by different colors). Size of scatter points is proportional to the number of trials for each combination, while continuous colored lines indicate the best-fitting performance of the Integrative model.

suggested by AIC values, the best model for reproducing human confidence choices was the Integrative one (Figure S5) which, as attested by lower RMSE values, better captured the observed performance in all three audiovisual conditions.

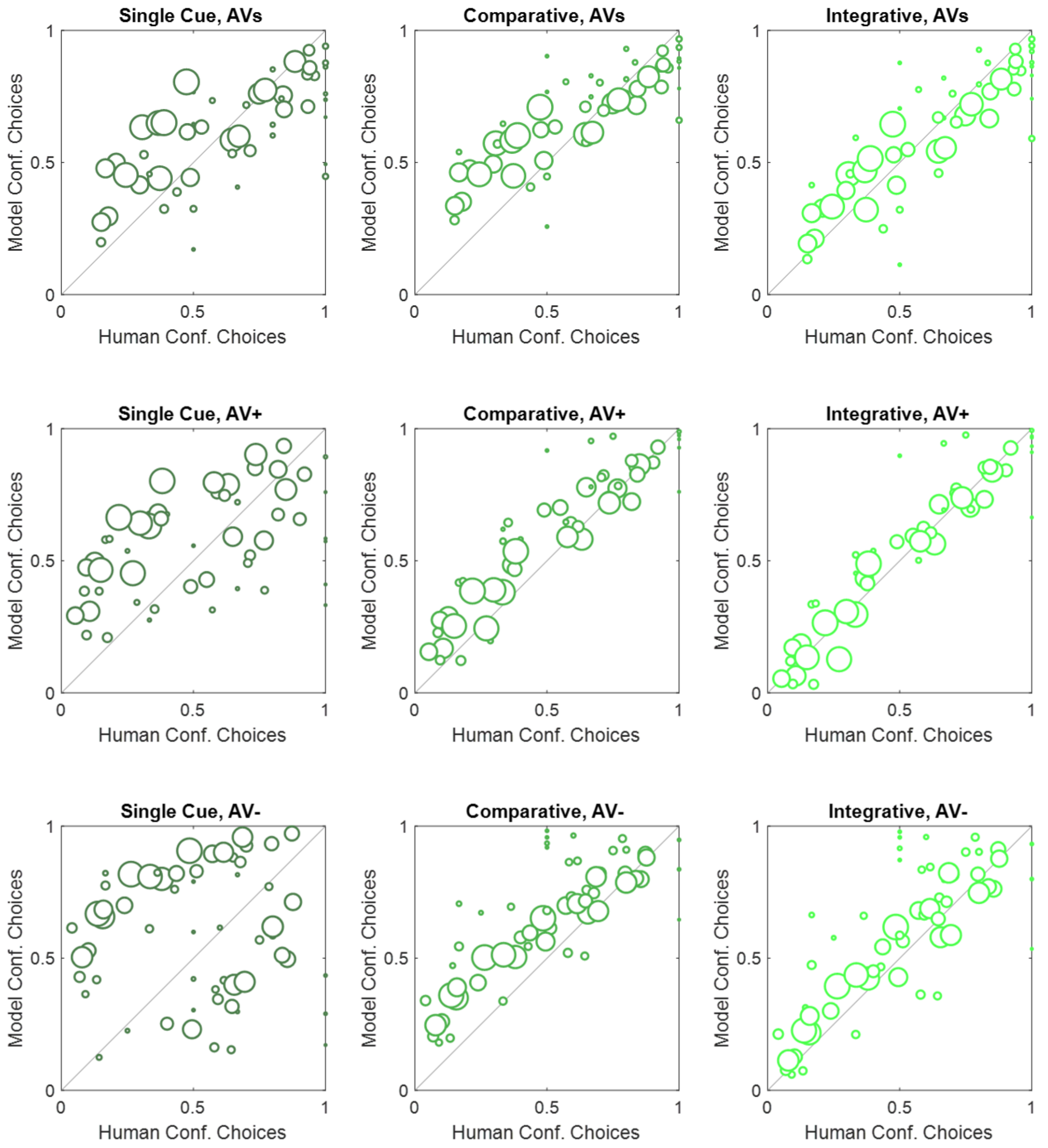


**Figure S6.** Goodness of fits of each model to human confidence choices for all bimodal conditions. Each point in each panel shows a confidence choice probability for a particular combination of stimulus difficulties across the two intervals and perceptual decision, for humans (x-axis) and the corresponding condition for the models (y-axis). Sizes are proportional to the corresponding number of trials performed by human participants for these conditions.

From the RMSE results, we further support our conclusion that the Integrative model performed better than the other two in terms of fidelity of the fits. As a final analysis of goodness of fit, we calculated the studentized residuals (point-by-point model minus human from Figure S6) for the Integrative model in all three bimodal conditions (Figure S7). We then conducted an outlier test by checking whether any of the residuals fell outside the ±3 range[5]. As no value exceeded this threshold, we could safely conclude that the Integrative model was a good fit of the human confidence choices measured in our study.

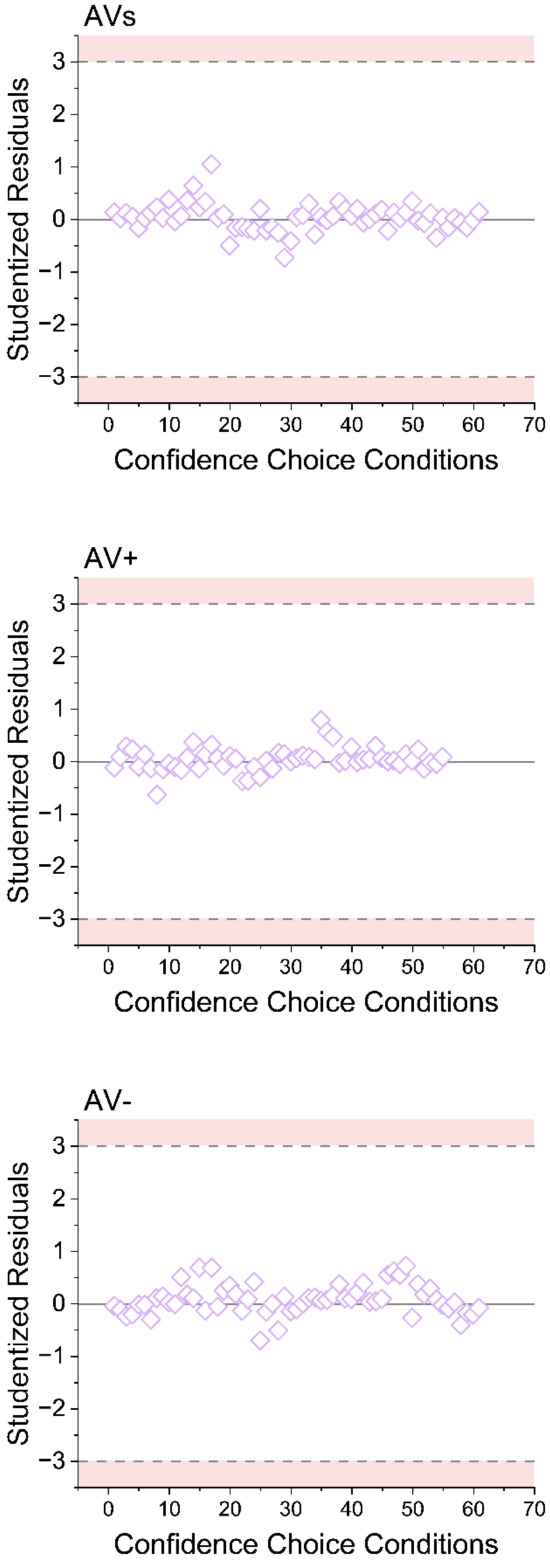


**Figure S7.** Studentized residuals for the regression models fitted between the Integrative model's confidence choice probabilities and the corresponding human choices. Red-shaded areas delimit the range outside which studentized residuals identify outliers, i.e., when the Integrative model would significantly misrepresent the human data.

| | AVs | | | AV+ | | | AV- | | |
|---|---|---|---|---|---|---|---|---|---|
| | SC | COMP | INT | SC | COMP | INT | SC | COMP | INT |
| RMSE | 0.194 | 0.172 | 0.114 | 0.265 | 0.107 | 0.069 | 0.386 | 0.159 | 0.108 |

**Table S2.** RMSE values computed between the human and model's performance.

### Details of the Metacognitive Models

#### Single-cue model

In Figure S8A, we show a schematic description of how metaperceptual evidence, for a given audiovisual interval, is generated with the Single-Cue model. At the perceptual level, visual ($S_V$) and auditory ($S_A$) sensory estimates for the visual ($\mu_V$) and auditory ($\mu_A$) physical intervals are obtained by corrupting these intervals with corresponding sensory noise ($\sigma_V$ and $\sigma_A$, respectively). Sensory estimates are modeled considering that observers have, on average, an unbiased internal representation of the physical interval (i.e., the mean of $S_V$ is $\mu_V$), as well as assuming both sensory noises to be normally distributed, so that $S_V \sim N(\mu_V, \sigma_V^2)$ and $S_A \sim N(\mu_A, \sigma_A^2)$.

At the perceptual level, both estimates are combined into a unitary bimodal percept following optimal multisensory integration[3]: each estimate is weighted according to its perceptual reliability $\omega_V$ and $\omega_A$, which are calculated as the inverse of the corresponding sensory variance and sum up to one. Thus, visual and auditory weights are computed as follow

$$\omega_V = \frac{\sigma_A^2}{\sigma_V^2+\sigma_A^2} \ , \qquad \text{(S6)}$$

$$\omega_A = \frac{\sigma_V^2}{\sigma_V^2+\sigma_A^2} \ . \qquad \text{(S7)}$$

The overall bimodal percept (P) can then be predicted to be

$$P = S_V\ \omega_V + S_A\ \omega_A \ . \qquad \text{(S8)}$$

The percept P is then compared to an internal sensory (bimodal) criterion $\theta_{AV}$, which is considered stable for each participant across the entire testing procedure and reflects interindividual differences in sensory reliabilities

$$\theta_{AV} = \theta_V\ \omega_V + \theta_A\ \omega_A \ , \qquad \text{(S9)}$$

where $\theta_V$ and $\theta_A$ are the unisensory visual and auditory criterion, respectively. For any given trial, if $P > \theta_{AV}$ participants will respond that S2 was closer in time to S3; otherwise, if $P < \theta_{AV}$, participants will respond that S2 was closer in time to S1. Crucially, the value obtained in equation S9 was used to model audiovisual performance for all bimodal conditions (AVs, AV+, and AV-), naturally assuming that the criterion did not change due to stimuli configurations. Rather, our models implied that changes in the BPs observed in the AV+ and AV- conditions reflected the shift in bimodal percepts conveyed by the audiovisual conflicts. To compute confidence in their perceptual decision, the Single-Cue model assumes that participants build some evidence $E_V$ from the visual cue alone, irrespective of the auditory one. Following this logic, $E_V$ is a combination of visual information used for the decision and a fraction of new information coming from the visual stimulus determined by the confidence boost parameter $\alpha_V$ (Mamassian & de Gardelle, 2022)

$$E_V = \alpha_V \, \mu_V + (1 - \alpha_V) S_V \ . \tag{S10}$$

The evidence $E_V$ is then converted into a distance to the sensory criterion $\theta_V$ and normalized relative to the sensory noise $\theta_V$, so as to obtain a unitless decision variable. Lastly, this evidence is corrupted by some confidence noise specific to the visual modality $\varepsilon_V$, so that equation S10 becomes

$$C_V = \frac{E_V - \theta_V}{\sigma_V} + \varepsilon_V \ . \tag{S11}$$

For the Single Cue model, the confidence evidence $C_V$ defined in equation S11 is generated similarly in the first and second interval of each confidence trial, even though the second interval contains a multisensory stimulus. The modulus of these confidence values is then compared across intervals, and the confidence judgment is computed such that the model will choose as more confident the perceptual decision bearing the higher absolute value for the confidence evidence.

For any given interval, we also compared $C_V$ with the original perceptual decision ($S_V > \theta_V$). If $S_V > \theta_V$ and at the same time $C_V > 0$, then the confidence evidence contrasted with the perceptual decision (as if participants realized they made a mistake in the perceptual task); when this happened, we replaced $C_V$ by $-|C_V|$. In doing so, we ensure that the model never selected these intervals when compared against any $C_V$ in agreement with the corresponding perceptual decision.

#### Comparative Model

A schematic representation of the comparative model, and how the confidence evidence $C_{AV}$ is computed from the audiovisual bisection judgment, is reported in Figure S8B. While the Comparative model does not differ from the Single-Cue one at the sensory level, it assumes that both visual and auditory confidence evidence $C_V$ and $C_A$ (with the latter being computed similarly to $C_V$, by applying equations S10 and S11 on the auditory performance) are generated independently and then combined according to the combinatory parameter $\beta$ as follows

$$C_{AV} = \beta \, C_V + (1 - \beta) \, C_A \ . \tag{S12}$$

For this computation, $\beta$ is a parameter free to vary and constrained between 0 and 1. Notably, this formulation allows us to test three possible scenarios: the two unisensory confidence evidence are combined according to the corresponding sensory reliabilities (i.e. $\beta = \omega_V$); the two unisensory confidence evidence are combined according to the corresponding confidence reliabilities (which we assume is the inverse of the corresponding confidence noises, i.e. $\beta = \frac{\varepsilon_A^2}{\varepsilon_V^2 + \varepsilon_A^2}$); or the bimodal confidence evidence is generated by simply considering the higher unisensory confidence evidence, so that either $\beta = 1$ (if $C_V > C_A$) or $\beta = 0$ (otherwise).

#### Integrative Model

Figure S8C illustrates the cascade of events that represents the Integrative model. In contrast to the previous ones, confidence evidence in the Integrative model is generated from the bimodal sensory percept $P$ (see equation S8), rather than from individual unisensory estimates. While the formulation at the sensory level follows once again similar principles, the evidence $E_{AV}$ is determined at first as

$$E_{AV} = (\beta_1\ \alpha_V\ \mu_V + (1-\beta_1)\ \alpha_A\ \mu_A) + (1-\beta_1\ \alpha_V - (1-\beta_1)\ \alpha_A)\ P\ , \tag{S13}$$

where $\alpha_V$ and $\alpha_A$ are the confidence boost for the visual and auditory modalities, respectively; $\mu_V$ and $\mu_A$ are the visual and auditory physical intervals, respectively; $P$ is the bimodal estimate; and $\beta_1$ is a free parameter that varies between 0 and 1, that determines the amount of sensory information coming from the visual rather than auditory modality to provide some confidence boost. The aim of equation S13 is to compute the amount of information used for the confidence evidence that includes some information that was not used for the sensory decision, and is the audiovisual generalization of equation S10. Confidence evidence is then converted and rescaled by the bimodal sensory sensitivity, before being corrupted by the bimodal confidence noise (similarly to what shown for the visual modality in equation S11)

$$C_{AV} = \frac{E-\theta_{AV}}{\sigma_{AV}} + \beta_2\ (\varepsilon_V + \varepsilon_A)\ , \tag{S14}$$

where $\theta_{AV}$ is calculated according to equation S9; $\varepsilon_V$ and $\varepsilon_A$ are the visual and auditory confidence noise, respectively; $\beta_2$ is a free parameter rescaling the sum of confidence noises; and $\sigma_{AV}$ is the bimodal sensory noise, which is calculated according to the optimal integration framework[3]

$$\sigma_{AV} = \sqrt{\frac{\sigma_V^2\ \sigma_A^2}{\sigma_V^2 + \sigma_A^2}}\ . \tag{S15}$$

Notably, the free parameter $\beta_2$ was introduced to avoid the direct prediction of how confidence noises were combined. Nonetheless, depending on its best value, we can still test different possibilities: if $\beta_2 < 1$, then audiovisual confidence noise is less than the sum of unisensory confidence noises; if $\beta_2 \sim 1$, then unisensory confidence noises are summed together; if $\beta_2 > 1$, then there is even more bimodal confidence noise than that predicted from the sum of unisensory confidence noises.

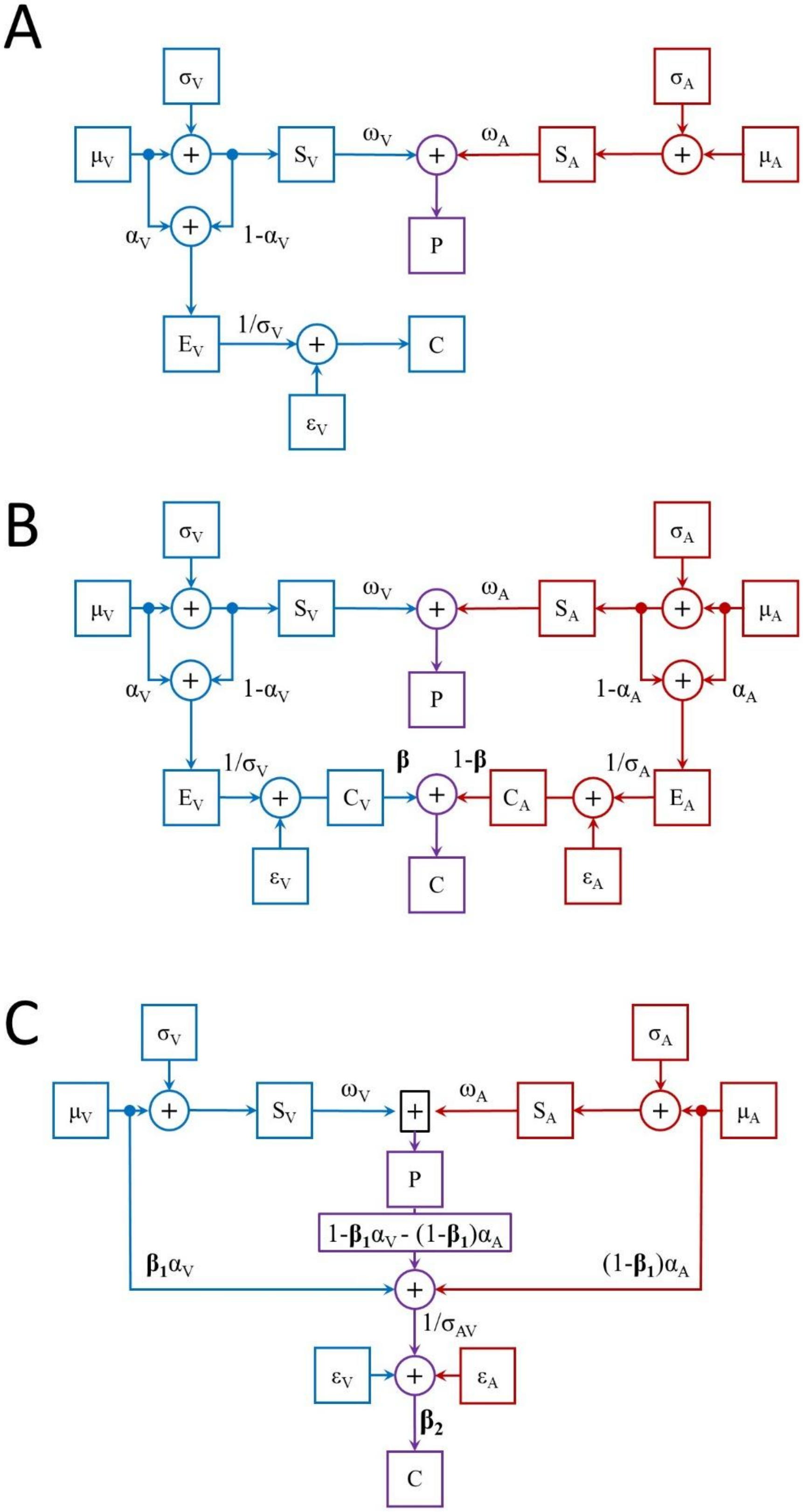


**Figure S8.** Graphical representations for the Single-cue (A), Comparative (B), and Integrative (C) models. Blue outlines indicate computations involving only the visual modality, red outlines identify computations involving only the auditory modality, and purple outlines highlight computations involving audiovisual integration.

#### Model Recovery

To further support our findings, we performed a model comparison analysis to demonstrate that outputs from the models defined for the current study could be easily distinguished. Starting from the empirical dataset, we thus simulated data from the Single Cue, the Comparative, and Integrative models in each bimodal condition (using four stimulus difficulty levels). We then generated 1000 datasets by randomizing, for each iteration, both sensory noises, sensory criteria, confidence boosts, and confidence noises. When there were free parameters, we randomized these parameters as well ($\beta$ for the Comparative model, $\beta_1$ and $\beta_2$ for the Integrative model).

Sensory noises were simulated starting from the empirical JNDs, randomly selecting values over $\left[\frac{\sigma}{2}, 2\sigma\right]$. Sensory criteria were selected over [0.4, 0.6] (as a reminder, the physical BP was fixed at 0.5), confidence boosts over [0, 1], and confidence noises over [0.25, 3] uniformly in log-space. For the comparative model, $\beta$ was simulated over [0, 1], while for the Integrative model $\beta_1$ over [0, 1] and $\beta_2$ over [0, 2]. Stimulus strengths were directly extracted from the empirical matrices and corresponded to [0.39, 0.46, 0.55, 0.62].

All simulated datasets were fitted by the three models and, for each iteration, we evaluated which model best represented the data from a maximum likelihood viewpoint. Overall, the simulations showed that the three models were easily recovered over the range of parameters tested (Figure S9). Across all three bimodal conditions, the Single Cue model was recovered 78% of the time, the Comparative model 95% of the time, and the Integrative model 99% of the time.

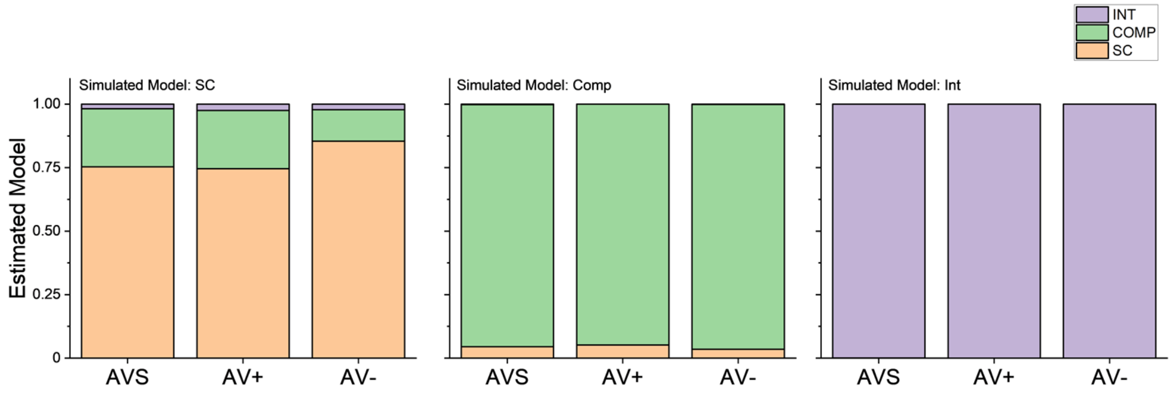


**Figure S9.** Results of the Model recovery analysis. Each panel reports the proportion of times each estimated model (SC, Comp, and Int) achieved the best fit.

Interestingly, when simulating data with the Single Cue model, sometimes the Comparative model was recovered (Figure S9, left panel). This is unsurprising, as the Single Cue model is nested into the Comparative (if $\beta = 1$, the two models are virtually identical). In light of this consideration, we further tested whether the two models were distinguishable at the empirical level by implementing a likelihood ratio test for nested models[6]. For this comparison, the test statistic corresponds to $\lambda_{LR} = -2(\lambda_1 - \lambda_2)$, where $\lambda_1$ is the log-likelihood of Model 1 and $\lambda_2$ is the log-likelihood of Model 2. If Model 1 is better at describing the empirical data, the test statistic is asymptotically distributed following $\chi^2$ (with the number of degrees of freedom that is equal to the difference of parameters' number, in our case, 1). Considering the log-likelihoods in our study $\lambda_{SC} = -6464$ and $\lambda_{COMP} = -5151$, the likelihood ratio test resulted in $\lambda_{LR} = 2626$ and $p < 0.001$. Given the statistical significance reported here, we could safely conclude that the two models differed from each other, even though the Single Cue was nested into the Comparative model.

#### Parameter Recovery

After demonstrating that all three models could be distinguished in their outputs, we tested how reliably confidence parameters could be estimated from randomly chosen values within the Integrative model, since this was the best model to describe our data. Starting from the datasets simulated in the previous section (which included 1000 simulations), we

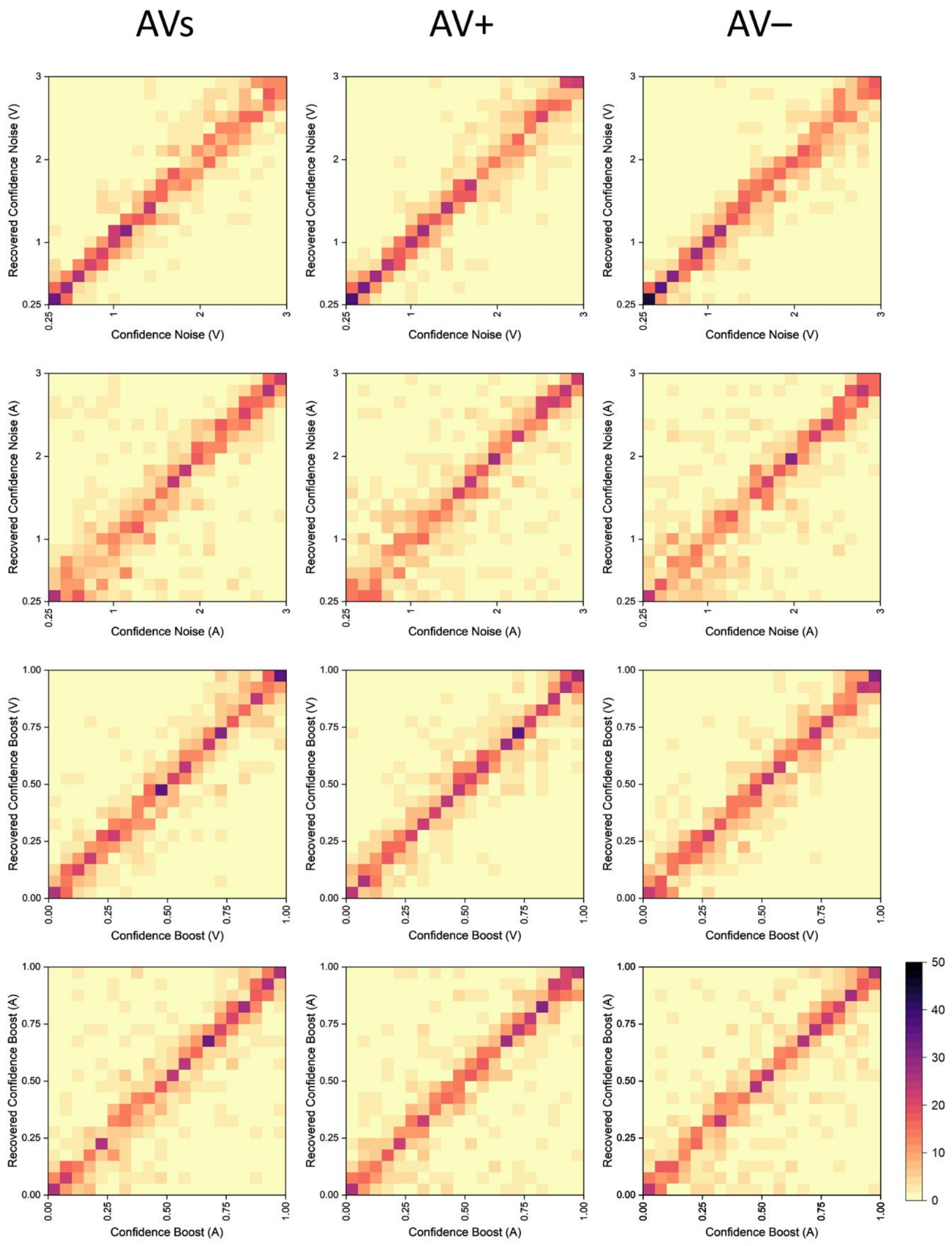


**Figure S10.** Parameter Recovery results for the Confidence Noise (V, first row; A, second row) and Confidence Boost (V, third row; A, fourth row). Each matrix shows the result of the parameter recovery performed on 1000 simulations, while each tile was created by dividing the corresponding parameter space into 20 equally-sized bins. The color of each tile represents the number of data points that are included in a given bin.

then recovered, for each iteration, one parameter from any of the confidence boosts, any of the confidence noises, and any of the free parameters ($\beta_1$ and $\beta_2$). As illustrated in Figure S10 and S11, all parameters were successfully recovered from the corresponding audiovisual condition. Overall, these findings further strengthened our implementation of the Integrative model, advocating in favor of its stability over a wide range of potential parameters.

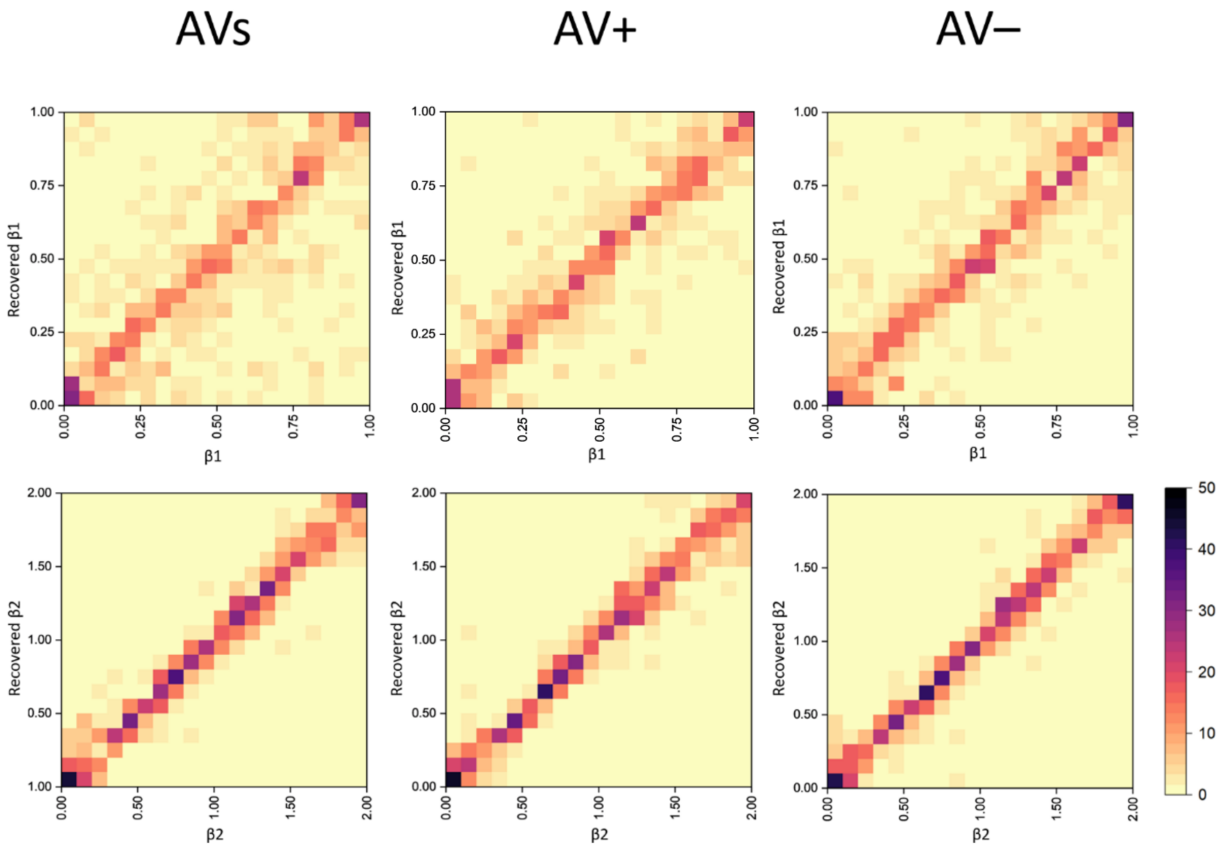


**Figure S11.** Parameter Recovery results for $\beta_1$ (first row) and $\beta_2$ (second row). Each matrix shows the result of the parameter recovery performed on 1000 simulations, while each tile was created by dividing the corresponding parameter space into 20 equally-sized bins. The color of each tile represents the number of data points that are included in a given bin.
